# Limited generalizability of dynamic fMRI correlates of adolescent rumination

**DOI:** 10.1101/2025.08.29.673124

**Authors:** Isaac N. Treves, Madelynn S. Park, Jamaal Spence, Nigel Jaffe, Kristina Pidvirny, Anna O. Tierney, Aaron K. Kucyi, John D.E. Gabrieli, Randy P. Auerbach, Christian A. Webb

**Affiliations:** McGovern Institute for Brain Research, Massachusetts Institute of Technology, Cambridge, MA; Department of Brain and Cognitive Sciences, Massachusetts Institute of Technology, Cambridge, MA; Department of Psychiatry, Columbia University, New York, NY, USA; Division of Child and Adolescent Psychiatry, New York State Psychiatric Institute, New York, NY, USA; Department of Psychiatry, Harvard Medical School, Harvard University; Center for Depression, Anxiety, and Stress Research, McLean Hospital, Belmont, MA; Department of Psychological & Brain Sciences, Drexel University, Philadelphia, PA

## Abstract

Rumination, or perseverative negative self-referential thinking, is a hallmark of depression. In adults, a dynamic resting-state fMRI model of trait rumination was recently identified through predictive modelling. In adolescents, a development period during which rumination and depression increase, the neurobiological correlates of ruminative thinking are less clear. In the current preregistered study, we examine dynamic connectivity correlates of self-reported rumination in the largest sample of adolescents to date (*n* = 443, containing clinical and non-clinical individuals). Notably, the adult model failed to generalize to our sample. In addition, linear models trained on default-mode network (DMN) connectivity, as well as whole-brain connectome models, failed to generalize to held-out data. In an exploratory random forest analysis, we found significant prediction performance of a model where increased variability between DMN-cerebellum, DMN-dorsal attention network, and DMN-DMN connections was nominally associated with higher rumination. However, the model did not generalize to an external sample with lower rumination scores and a distinct scanner protocol. Our findings illustrate the difficulty of characterizing the neurodevelopment of risk factors for depression.

## Main Text

Rumination, or perseverating on negative self-referential thoughts, memories, and one’s own negative mood, is present across a range of mental health disorders.^1,2^ Rumination is a transdiagnostic predictor of feelings of hopelessness,^3^ psychological distress,^4^ and suicidal ideation,^5^ and levels of ruminative thinking may be highest in adolescents and adults with co-occurring anxiety and depression.^6,7^ For this reason, rumination may be a valuable treatment target for the prevention of these disorders. As many anxiety and depressive disorders first emerge in adolescence,^8^ there is a clear and present need to understand how ruminative thinking arises in adolescents. One step toward this goal is to identify individual differences in rumination and their biological bases in the brain.

To study individual differences in rumination, researchers have typically relied on retrospective self-report questionnaires. In adults, questionnaires like the Ruminative Response Scale decompose rumination into subscales of brooding, reflection, and depressive rumination.^9^ In adolescents, similar constructs are measured using scales like the Children’s Response Style Questionnaire.^10^ These scales inquire about individuals’ tendencies to ruminate, and are strongly correlated with depression.^10,11^ Recent work has highlighted the value of state-level factors using ecological momentary assessment (EMA).

EMA minimizes recall biases by asking what individuals experience in the moment. Studies utilizing EMA to investigate rumination highlight its links to stress and negative affect in daily life. Momentary rumination predicts subsequent negative affect (and vice versa),^12^ and rumination also mediates the role of life stress and negative events in prospectively predicting negative affect and depressive symptoms over several weeks in both clinical^13^ and nonclinical^12,14^ samples.

In parallel, researchers have studied the brain bases of depressive rumination. Among adolescents and adults, fMRI research highlights increased activation during rumination within the default-mode network (DMN),^15^ an intrinsic functional brain network associated with self-referential processing and internally-focused thought.^16,17^ More broadly, the DMN is activated not only during rumination, but when attention is directed away from external stimuli and more internally^18^; for example, during mind-wandering^19^ and autobiographical memory retrieval.^20^ This work suggests that ruminating individuals may be more likely to focus on internal mental states and autobiographical information (especially negatively-valenced content) relative to individuals low in rumination.^15^ Increased DMN and subgenual prefrontal cortex (sgPFC) activation during rumination versus control conditions^15^ aligns with a proposed model that indicates increased functional connectivity between the DMN and sgPFC that accounts for increased rumination in MDD.^21^ Further, activation in the amygdala and subgenual anterior cingulate (sgACC),^22^ as well as the posterior cingulate cortex (PCC), may be heightened during ruminative states.^22–24^

### Brain Correlates of Individual Differences in Rumination

Despite this informative work, it is unclear what brain differences are present in individuals who tend to ruminate. There is not a clear activation—or pairwise connectivity—difference that reliably predicts individual differences in depressive rumination. For example, a study with over a hundred unmedicated adults and healthy controls found no correlation between static DMN connectivity and trait rumination.^25^ However, a multivariate fMRI study of trait rumination in adults indicated that dynamic, or time-varying, connectivity is a promising marker of the phenotype.^26^

In the study, time-varying connectivity between the DMPFC (a node of the DMN) and the rest of the brain predicted individuals who self-report ruminative brooding (hereafter referred to as the DMPFC-Rum model).^26^ This finding, validated across multiple samples, is notable because it implicates a specific area of the DMN—the DMPFC—which could be a possible treatment target. Neuromodulatory interventions for depression often involve transiently disrupting activity in the prefrontal cortices, including the DMPFC.^27^ In addition, this work aligns with other evidence suggesting that dynamic connectivity is especially sensitive to individual differences^28–30^, and work indicating that rumination has been associated with decreased temporal stability in DMN regions.^31^ Perhaps dynamic connectivity captures state-level differences in cognition—specifically, the repetitive, constrained negative states that may emerge during induced rumination or spontaneously during resting states.

In adolescents, there are no well-validated neural markers of rumination, and most work has been conducted in small sample sizes (n∼50).^32,33^ Some research has identified specific brain regions linked with rumination in depressed adolescents. Notably, reduced resting-state functional connectivity between the sgACC and both the inferior frontal gyrus (IFG) and middle frontal gyrus (MFG) – key regions comprising the central executive network responsible for attention and high-level cognitive functioning^34^ – has been associated with elevated levels of rumination.^35^ Other investigations have found that abnormal frontoinsular-DMN connectivity may contribute to maladaptive ruminative tendencies in depressed adolescents.^33^ Research has also highlighted aberrant DMN connectivity, particularly between the PCC and sgPFC, as being positively correlated with trait rumination.^32^

Further, a recent longitudinal resting-state fMRI study of adolescents transitioning into adulthood found that maladaptive ruminative processes—brooding and depressive rumination—are associated with changes in static functional connectivity, including increased connectivity in the DMN and right frontoparietal network (FPN), and reduced connectivity within the salience and limbic networks.^36^ Notably, static connectivity within the salience network longitudinally mediated the relationship between brooding and internalizing symptoms. Dynamic functional connectivity changes were observed but were less robust, with some evidence that DMN dynamics—particularly in the superior middle frontal gyrus— mediated the relationship between depressive rumination and externalizing symptoms.^36^ Despite these findings, resting-state markers of rumination remain preliminary, constrained by small sample sizes and heterogeneous methodologies.

### Present Study

Methodologically rigorous, large sample studies are needed to assess brain markers of rumination in adolescents. This work could help identify the biological bases of depression and provide treatment targets. This is part of the ‘reciprocal validation’ model of psychiatric neuroimaging,^37^ where large sample correlational findings (at single timepoints) could inform smaller-sample treatment/longitudinal studies.

In keeping with this approach, we conducted the first large-scale, multi-site study on fMRI correlates of adolescent rumination (**Fig. 1, Supplementary Tables 1-5**). Four cohorts were collected at the same site using identical scanners and included adolescents of similar age (*n* = 344); a fifth, independently collected cohort (*n* = 99), composed of younger and less ruminative adolescents, served as an external validation sample. Building on prior work demonstrating the predictive utility of the DMPFC-Rum model in adults,²□ we focused on dynamic resting-state connectivity (**Fig. 2**). However, given the substantial developmental changes in adolescent brain structure and function,³□ we also tested adolescent-specific models. We measured rumination using experience sampling probes (i.e., EMAs) and retrospective trait measures to compare momentary vs retrospective measures. We conducted four machine learning analyses based on understanding DMN and whole-brain relationships with rumination (**Fig. 3**):

1. (Preregistered) Tested the DMPFC-Rum model.^26^
2. (Preregistered) Trained our own models using other DMN seeds.
3. (Preregistered) Trained whole-brain connectivity models using connectome predictive modeling.^39^
4. (Exploratory) Trained DMN-based and whole-brain random forest regression models.

**Figure 1.**
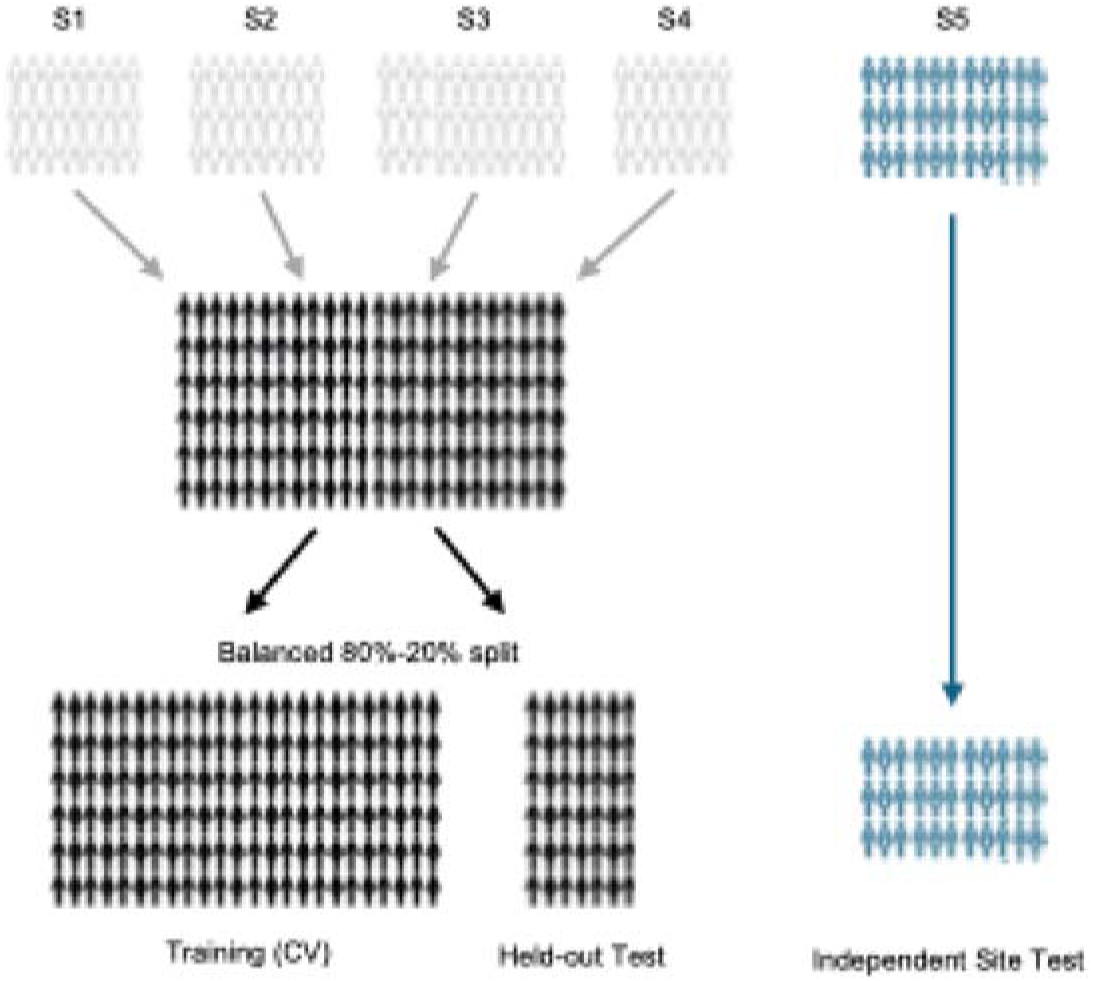
Predictive modelling approach. We leveraged 5 separate cohorts encompassing 486 adolescents. Four cohorts were collected by the same group and involved similar scanning protocols and populations. These cohorts were combined, shuffled, and then separated into a balanced 80% training set and 20% validation set. To test generalizability to external data, we then further assessed the performance of the models on the fifth cohort.

**Figure 2.**
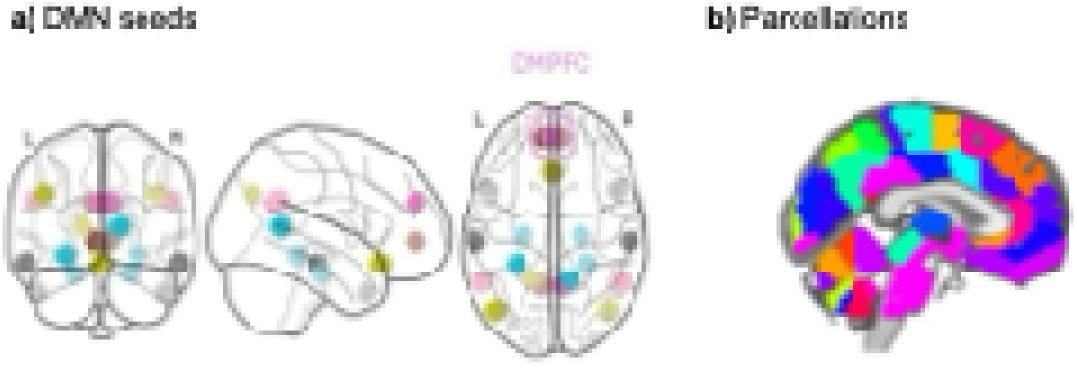
The seeds and ROIs used to construct dynamic connectivity features. **a**) 20 DMN seeds were used. Circled in pink is the dorsomedial prefrontal cortex (DMPFC). **b**) The Brainnetome parcellation with 280 regions was used.

**Figure 3.**
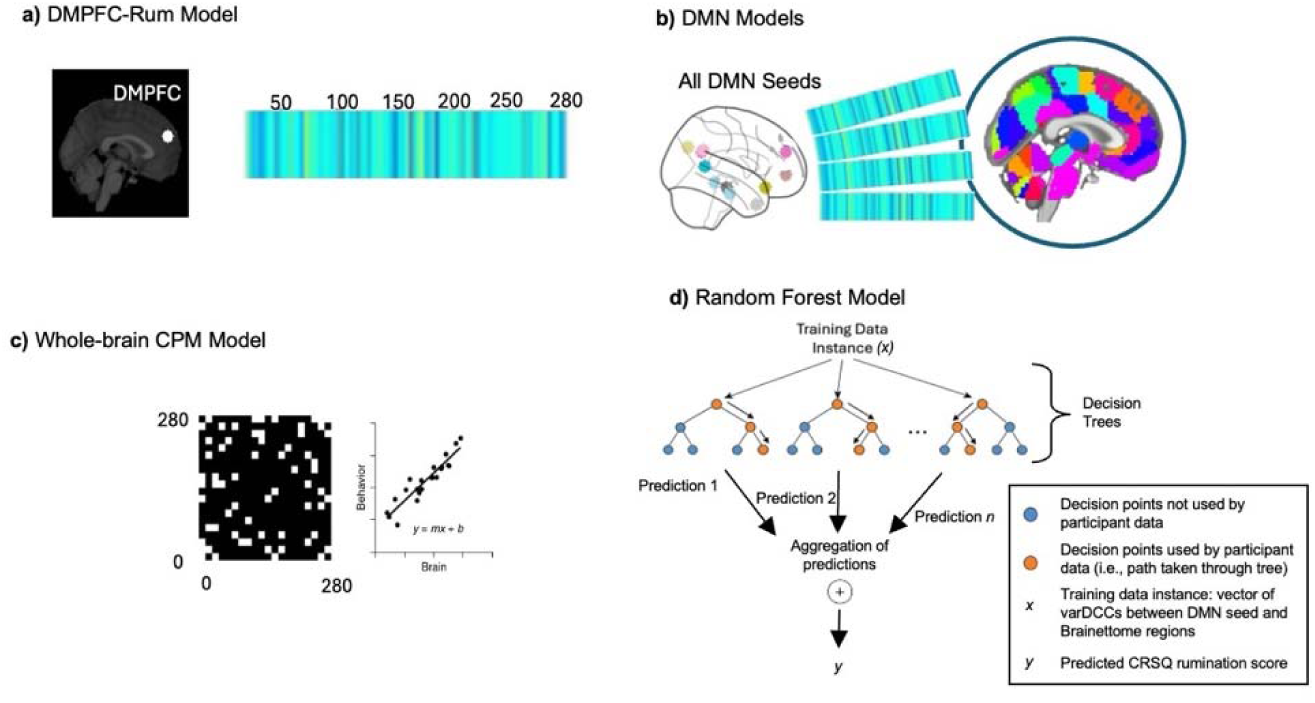
Modeling approaches. **a)** The DMPFC-Rum model, consisting of a set of beta weights (yellow is positive, blue is negative) over the 280 Brainnetome regions. **b)** We use CPM models on pairwise interactions between all 280 Brainnetome regions. **c)** We train our own models using lasso regression between the 20 DMN seeds and the 280 Brainnetome regions. **d)** We use random forest regression for each of the 20 DMN seeds and 280 Brainnetome regions. Note that we also applied random forest using all pairwise interactions between the 280 regions, which chiefly involves expanding the scope of the features *x*.

Our modelling approach involved rigorous techniques in brain-behavior analyses. We trained models, validated them in held-out data, and then tested valid models in external data, allowing for accurate characterizations of predictive generalizability.^40^ Due to relationships between rumination and rumination-related brain circuitry and factors like sex, clinical status, parental risk, and puberty (see **Supplementary Note 1**), we controlled for these factors. We examined multivariate models, which may capture the distributed networks that support adolescent cognition. Finally, as our dataset contains not only neurotypical but also clinical groups, there was sufficient variation in rumination scores to train robust models.

## Results

### Approach 1: Lack of predictive validity of adult-derived DMPFC-Rum model

Correlations were assessed between the model predictions and self-reported rumination scores (trait CRSQ, **Extended Data** Fig. 1, and EMA, **Supplementary** Fig. 1) across **Studies 1-4** *(n* = 344). There were no significant correlations between predictions and reported CRSQ (*r*(328) = .044, *p* = .43), nor EMA (*r*(326) = - .039, *p* = .48). There was no correlation between predictions and total puberty scores (*r*(342) = - .03, *p* = .58). When separating the predictions by dataset and clinical characteristics, there were no significant correlations between CRSQ rumination and model predictions (**Supplementary Table 6**). There was a significant positive correlation between EMA rumination and the predictions in the **Study 4** dataset (*r*(66) = .24, *p* = .045), and an inverse association in the **Study 3** healthy controls (*r*(62) = - .28, *p* = .023), neither were significant when controlling for multiple comparisons (**Supplementary Table 7**). No other subsets showed significant correlations for EMA. Scatter plots are shown in **Extended Data** Fig. 2 and 3.

### Approach 2: Lack of predictive validity of DMN seed models

Models based on connectivity between DMN seeds and the rest of the brain were trained using lasso regression. In the training set (*n* = 275), we found one CRSQ model that passed our selection threshold, from the right parahippocampal cortex (non-parametric *p* = .031). However, this model failed to predict scores in the test set (*n* = 69), (*r*(67) = - .16, *p* = .19). No EMA models passed our selection threshold. Full training set p-values are reported in **Supplementary Tables 8and 9**.

### Approach 3: Lack of predictive validity of whole-brain CPM models

Models were trained based on whole-brain connectivity matrices using CPM which finds negative predictive and positive predictive networks. In the training set (*n* = 275), we found that a negative network model for the CRSQ passed our selection threshold (*p* = .033), which was still significant when regressing out head motion (*ps* = .025) as well as puberty and head motion (*ps* = .004). However, the model did not generalize to the test dataset when conducting Pearson’s correlations (negative network; *r*(67) = - .048, *p* = .7). No models passed our selection threshold for predicting EMA scores, nor for either rumination measure when using static (averaged) connectivity (**Supplementary Table 10**).

### Exploratory Analysis: Partial generalization of Random Forest models

Models were trained based on DMN-seed and whole-brain connectivity, using random forest (nonlinear) machine learning models. In the training set (*n* = 275), 16 models passed selection threshold for the CRSQ (**Supplementary Table 11**). Only the Right and Left Temporal Pole models generalized to the test datasets (*r*(67) = .24, *p* = .05 and *r*(67) = .26, *p* = .032). The Right Temporal Pole was chosen for subsequent inspection given findings from external validation. An example regression tree (of 250) is shown in **Supplementary** Fig. 2. Importance maps over regions are shown in **Fig. 4**. The Right Temporal Pole model had high positive weights on cerebellum, precuneus, and temporal lobe (**Table 1A**) and negative weights on temporal gyri, hippocampus and cingulate (**Table 1B)**. The model (**Fig. 4b)** showed positive importances for the dorsal attention network (*p* = .013, FDR-*p* = .068), the default-mode network (*p* = .014, FDR-*p* = .068), and the cerebellum (*p* = .034, FDR-*p* = .11), suggesting dynamic connectivity with these regions positively predicted rumination (importance values are reported in **Supplementary Table 12**). Network properties of the temporal poles suggest they were less hub-like and connected in our sample (**Supplementary Note 2)**. We did not observe generalizable models of EMA rumination (**Supplementary Table 13**).

**Figure 4.**
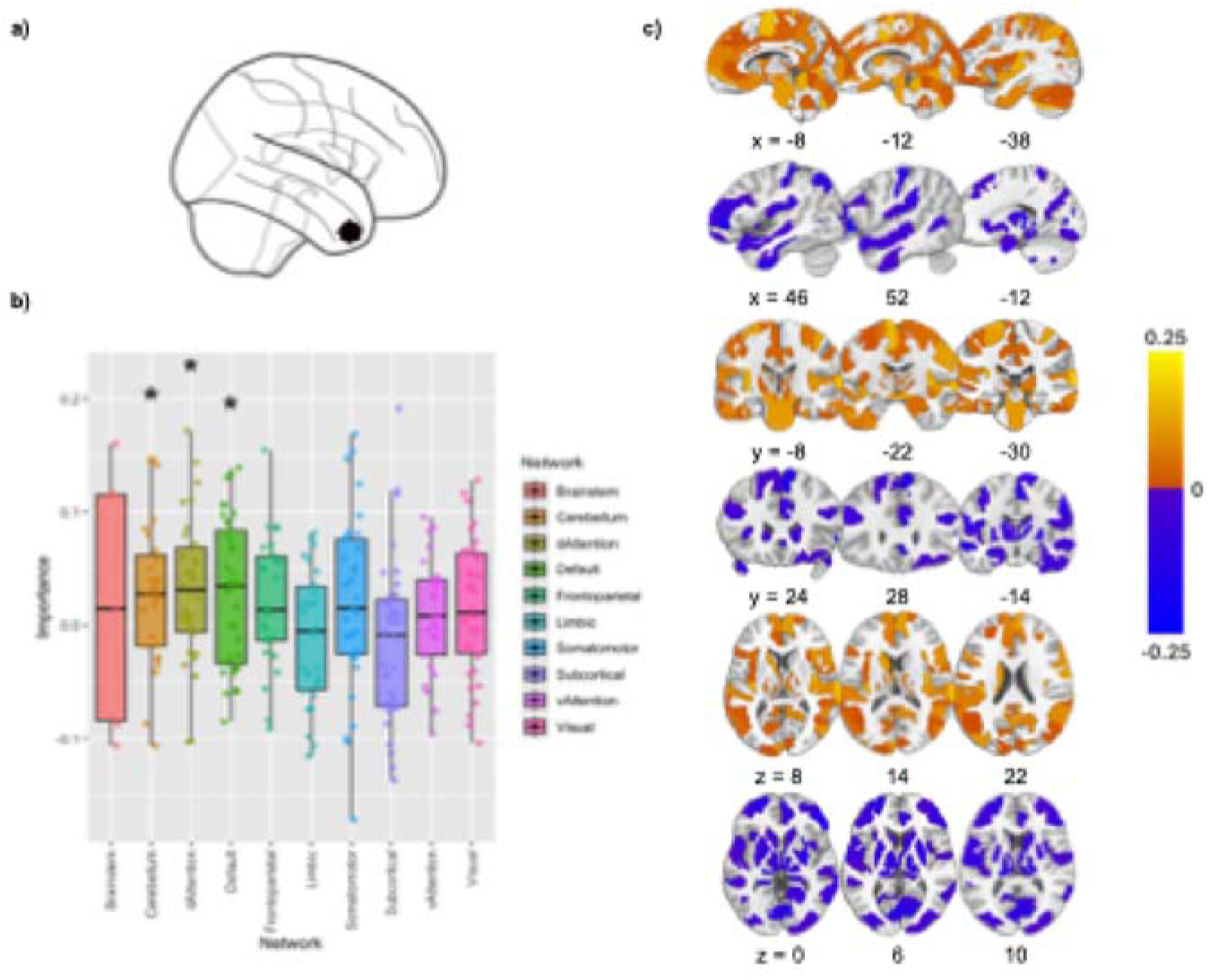
Right temporal pole seed and importance brain maps. **a)** Right temporal pole seed that was used in our exploratory analysis. **b)** Right Temporal Pole model, representation in terms of networks. Average importances and standard errors for each of the ten Brainnetome networks. **c)** Sagittal, coronal, and axial views of the greatest importance weights for the brain regions that displayed positive predictive weights (warm) and negative predictive weights (cool). The color bar represents the sign and magnitude of the weights. *, *p–*uncorrected < 0.05, two-tailed *t-*test compared to zero. The *p-values* are: dorsal attention network (*p* = .013, FDR-*p* = .068), default-mode network (*p* = .014, FDR-*p* = .068), and the cerebellum (*p* = .034, FDR-*p* = .11).

**Table 1.**
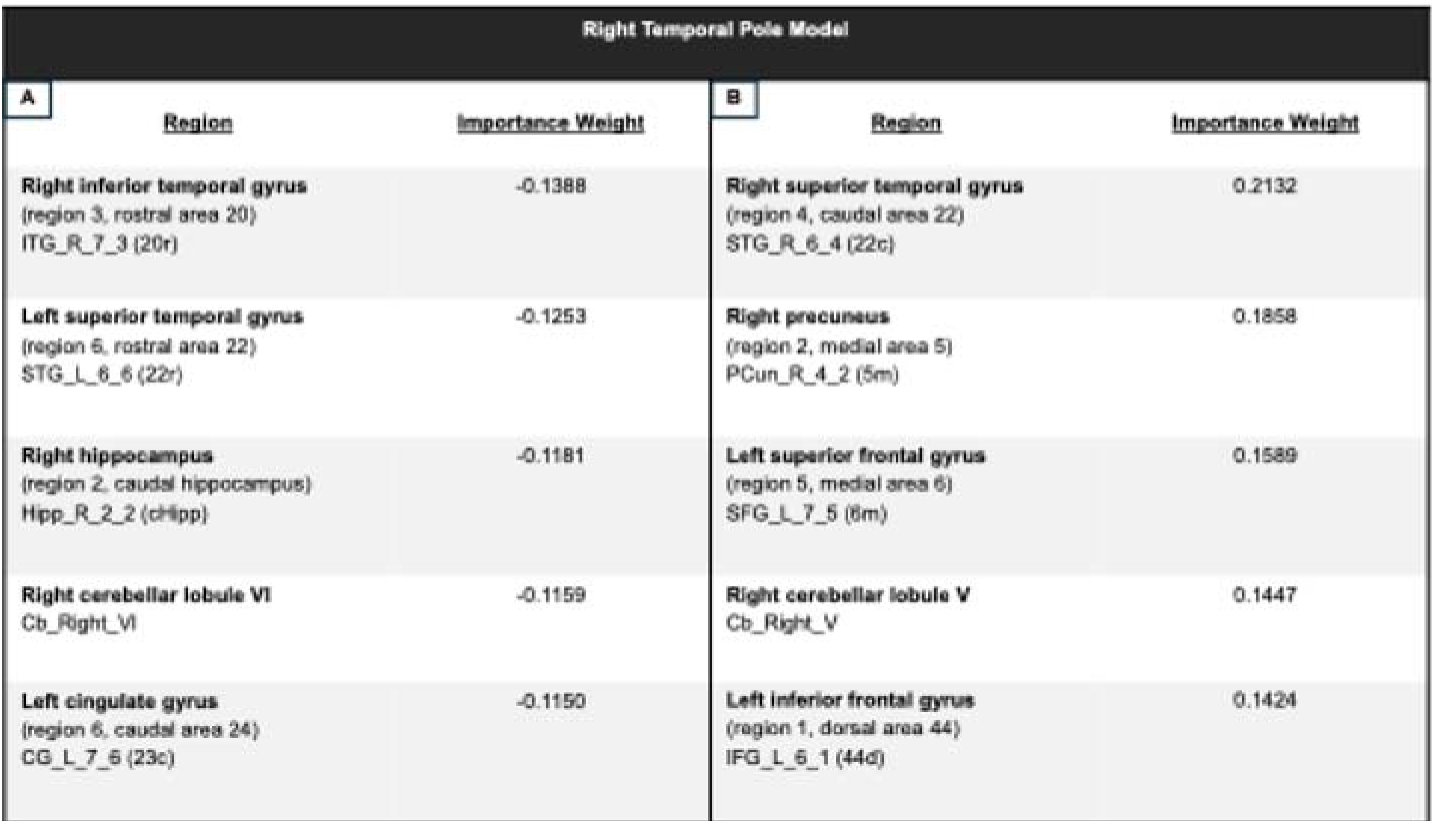
The table lists the (A) five greatest negative importance weights and the (B) five greatest positive importance weights and their associated brain regions for the Right Temporal Pole model. Brain regions were identified using the Brainnetome Atlas^48^ (BNA; https://scalablebrainatlas.incf.org/human/BNA).

In addition, a whole-brain random forest model using variance of DCC passed the selection threshold (*p* = .012) and trended towards significance in the validation dataset (*r*(67) = .24, *p* = 0.054). The model heavily implicated dorsal and ventral attention networks, as well as subcortical and cerebellum regions (**Supplementary** Fig. 3). No whole-brain models passed selection threshold for EMA (**Supplementary Table 14)**.

Interestingly, the Right Temporal Pole model showed an interaction with puberty where it performed better for higher puberty individuals (*B* = 0.82, *p* = .008, **Extended Data** Fig. 4). This was also found for the whole-brain model interaction (*B* = 0.33, *p* = .019).

### External Validation

The external test dataset (*n =* 99) was harmonized using COMBAT, and then used to evaluate the selected random forest models (Left Temporal Pole, Right Temporal Pole, whole-brain). None were significant when examining the full distribution of rumination scores (*p*s > 0.35, **Supplementary Table 15**), and there were no interactions with puberty (*p*s > 0.2). However, when accounting for lower CRSQ scores in the held-out dataset (**Supplementary** Fig. 4), we found that the Right Temporal Pole model showed a positive relationship that was similar in magnitude to the validation dataset (*r*(49) = 0.26, *p* = 0.064) (**Fig. 5)**. The Right Temporal Pole model also showed a significant negative relationship when not harmonizing using COMBAT (*r*(97) = −0.23, *p =* 0.02); the predictive relationship was reversed. For full external validity analyses including non-harmonized results see **Supplementary Note 3.**

**Figure 5.**
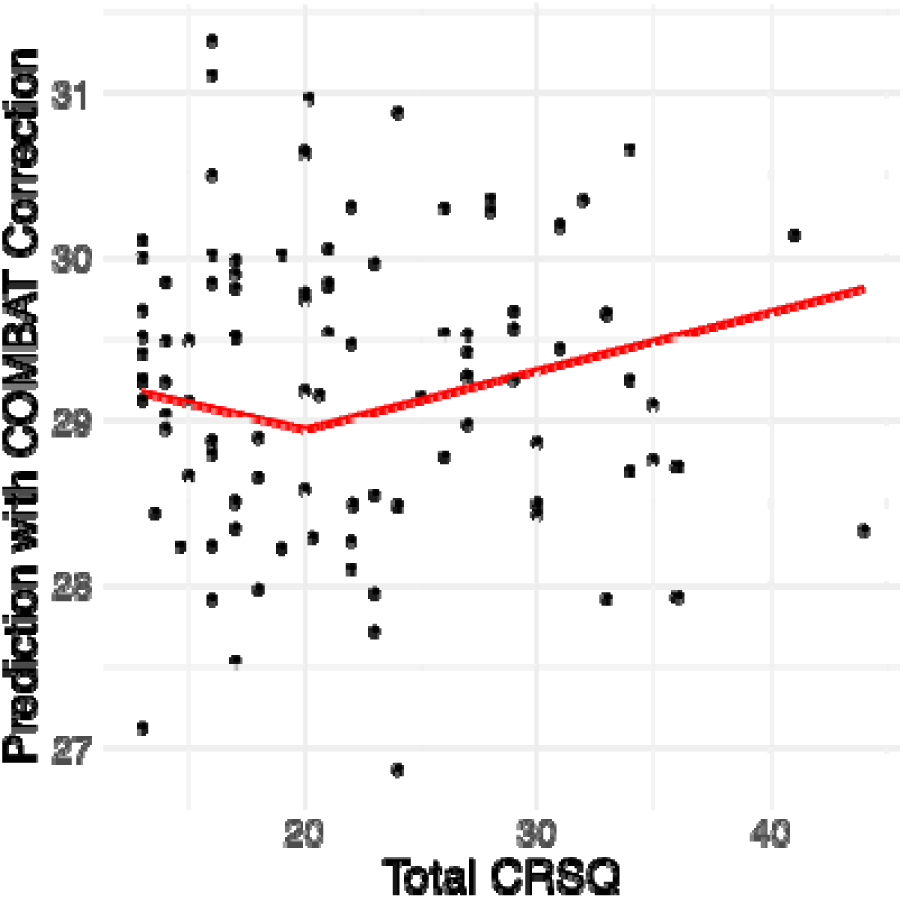
Prediction performance in harmonized external sample, piecewise fit. Red lines reflect piecewise linear regressions, with cutoff at CRSQ=20 (1 S.D. below mean of training set).

## Discussion

Rumination consists of perseverative negative self-referential thinking and is a hallmark symptom of depression, a disorder with marked increases during the adolescent years. Testing the link between rumination and brain measures could help inform brain targets for neuromodulatory treatment and potentially provide objective measures of depression risk and severity. There are concerns about the replicability of brain-behavior associations, and the following steps were taken to address them: (1) analyzed >400 adolescents, the largest sample in a neuroimaging study of trait rumination; (2) preregistered our main analyses; (3) trained multivariate instead of univariate models; (4) examined whether models generalized to external data, and (5) included participants with a spectrum of clinical symptoms. Despite this rigorous approach, none of our preregistered models generalized to the held-out data. While we did find suggestive results for an exploratory nonlinear model, our overall results highlight the difficulty of finding replicable brain-behavior correlates and challenge the feasibility of psychiatric neuroimaging.

We sought to predict trait rumination (as measured by the CRSQ^10^) and EMA rumination. These two operationalizations of rumination may capture different components. Whereas the CRSQ captures individuals’ narratives about themselves based on retrospective self-report (e.g., “I tend to ruminate”), EMA captures momentary responses averaged over time. To predict individual scores on these two measures, we generated dynamic functional connectivity from resting-state fMRI. We chose dynamic connectivity for two reasons: 1) it yielded generalizable correlates in adults^26^ and 2) dynamic connectivity may be more sensitive to self-report than static connectivity because it captures fluctuating cognitive states.^28,29^ Accordingly, we calculated the variance of dynamic conditional correlations between a set of 280 whole-brain regions as well as between those regions and 20 seeds in the DMN. Variance of DCC captures how two areas fluctuate over the course of a scan. We extracted DCC and rumination scores from five studies across three different resting-state protocols and two different scanners.

Our first approach was to test the adult DMPFC-Rum model^26^, consisting of 83 weights between the DMPFC (a DMN seed) and other brain regions (with greatest importance assigned to inferior frontal gyrus, cerebellum, and inferior temporal gyrus). This model generalized across three samples in a previous study, but it should be noted that it did not generalize to additional MDD samples.^26^ The model did not predict rumination in our adolescent samples as well, either when examining all individuals or each site individually. There are a number of reasons why the model could have failed to generalize including scanner differences, developmental differences, and rumination scale differences. Thus, we trained our own models using preregistered approaches.

In keeping with theories that the DMN is involved in rumination,^20^ we trained 20 models, each consisting of lasso-derived weights across 280 DCC features from each seed. We ensured that the training (80%) and test datasets (20%) were balanced across age, sex, clinical features, site, and puberty (as these can all have effects on rumination and brain measures, **Supplementary Text**). While some models passed selection in the training set (based on permutation testing), none generalized to the held-out data. After observing the failure of DMN-seeded models to generalize, we decided to assess whether brain regions outside the DMN could predict rumination. To test this, we trained connectome models on the whole-brain connectivity.^39^ Again, there was a failure to generalize.

### Limitations of preregistered approach

One possibility for the lack of generalizability is that even larger samples are required. One study^51^ examined univariate and multivariate models of behavior in the ABCD dataset (adolescents) and found that sample sizes in the thousands were required to uncover effect sizes with low variability (high replicability). Largest effect sizes were observed for multivariate models, rsFC, and cognitive outcomes. It should be noted, however, that there is some debate over whether multivariate models provide more replicable associations.^52,53^

Another possibility is that our sample contained too much phenotype variability. We controlled discovery and validation samples across age, sex, site, clinical subtype, and puberty. However, by blending across these factors we may have obscured true signal. For example, it may be the case that biological substrates of depression symptoms are distinct in those who are predisposed to recurrent depression (genetically or otherwise) vs those who experience a transient depressive episode triggered by a stressful life event and who have no underlying vulnerability to the disorder.^54,55^ In addition, results from exploratory models indicate that models perform better for individuals reporting more advanced pubertal development. Subsample analyses and analyses that are robust to overall developmental differences in functional brain organization could be useful in the future.^56,57^

There are a number of other possibilities. The resting-state fMRI scans may have been too short to find trait-like, reliable features and there may have been nonlinear effects of head motion not accounted for by our methods ^58^. We extracted timecourses from group averaged regions. Perhaps personalization of brain regions or networks could have led to more informative signals.^59^ Possibly, the addition of other features above and beyond DCC (e.g. static connectivity, task-based measures, or even EEG measures) could have improved predictive performance. Finally, both the lasso and CPM approaches regularize features by discarding non-associated variables, and both consist of linear combinations of features (with different weights for lasso, and binary (1, −1) for CPM). It is possible that there was meaningful information in the discarded features that could be captured in a nonlinear way. In addition, the rumination scores (trait and EMA) deviated from normality.

### Exploratory models show partial generalization

To test the importance of nonlinear relationships, we conducted an exploratory random forest regression analysis. Random forest regression consists of training a large number of decision trees (which are capable of uncovering non-linear relations) and averaging across their outputs to predict the outcome. Random forest models provide flexibility over interpretability.^60^ Previous studies have found random forest may be effective for predicting autism diagnoses,^41^ alcohol use disorders,^43^ and predicting cognitive outcomes.^42^ Interestingly, when compared directly to other machine learning models in neuroimaging applications, there is not consistent evidence that RF is better than other linear models in terms of reliability and prediction accuracy.^61,62^ However, in our sample, RF models generalized to predict CRSQ in internal and external held-out data where linear models such as CPM did not.

The most promising model incorporated connectivity between the Right Temporal Pole and the rest of the brain. The temporal poles are critical seeds within the DMN that integrate semantic, emotional, and autobiographical information.^63^ The left temporal pole plays a key role in linking high-level sensory representations with semantic information and conceptual labeling (e.g., naming or categorizing recognized stimuli), while the right temporal pole is involved in associating sensory inputs with emotional responses and socially relevant memory.^64,65^ For the Right Temporal Pole model, higher variability with the DMN, dorsal attention network (DAN), and cerebellum predicted higher rumination scores. This pattern indicates that trait rumination involves not only the specialized functions of the temporal poles, but also their dynamic coupling with the networks supporting self-referential thought (DMN), attentional control (DAN), and sensorimotor processes (cerebellum). Increased variability of these connections may suggest neural mechanisms underlying the internally focused and inflexible thought patterns that are characteristic of rumination, consistent with prior work associating altered DMN and DAN connectivity to maladaptive self-focus and difficulties in attentional disengagement in rumination.^21,66^

In our data, the right temporal pole model not only predicted CRSQ in the validation data (data similar to the training set but not exposed to the model) but also showed partial generalization in distinct external data. Specifically, the model showed a positive trend relationship with CRSQ scores in adolescents who were similar in rumination to the training and validation datasets. There are a couple reasons why we believe this non-significant finding deserves consideration. First, the external dataset may have included more resilient individuals as it excluded depressed and anxious adolescents, resulting in much lower CRSQ scores. Thus we believe the subsample approach is reasonable. Second, when not harmonizing connectivity (input features to the model), the Right Temporal Pole model showed a significant *negative* relationship with CRSQ, suggesting that it contains information relevant to the rumination but depends on scanner features. Third, the adult DMPFC-Rum model also linked increased variability between DMN and DAN to higher rumination scores (as well as DMN-IFG variability).

Finally, a recent mega-analysis of 400 youth MDD individuals and 400 controls implicated higher static connectivity in the DAN and lower connectivity in the DMN with diagnosis and symptom severity.^67^ Together, these converging findings suggest the hub-like networks of the DAN and DMN deserve particular highlighting as possible mediators of disrupted introspective attention. Of course, the suitability of the current model for large-scale application as a biomarker is inadequate given sensitivity to scanner characteristics, rumination score distributions, and pubertal development.

### Future directions

The current cross-sectional approach is limited in understanding the developmental trajectory of rumination or how brain signatures may differ by age group. One potentially promising avenue for future research is to examine random forest brain prediction models in longitudinal/treatment studies. However, it is unclear why the random forest model performed better than other models, and why trait but not EMA rumination was predictable. Perhaps, like others have found, sparse models (like lasso) perform worse.^62^ As mentioned, the temporal pole models that generalized did not pass traditional significance testing in the external data. Thus, the model should be considered very preliminary, and further investigation of the implicated networks and regions (DMN, DAN, cerebellum) is a necessity.

It is clear that work needs to be done examining the tradeoffs between different multivariate models including deep learning methods^68^ in prediction and classification applications. The different models should be characterized in terms of reliability, prediction accuracy, and feature relevance. It is plausible that effective models vary on the data and the outcome. It is also increasingly clear that feature weights from models implicate wide ranges of brain areas, outside of canonical networks like the DMN.^69^

Finally, research on brain-behavior correlations may need to be assayed on a population level (Ns > 1000) like genetic association studies. This severely limits their clinical utility, based on cost, small effect sizes, and specificity. A recent study leveraged multimodal neuroimaging data to classify individual diagnosis of depression, training over a million models in a discovery sample of 1000 participants.^70^ The best held-out accuracy that they could achieve was 62%. For comparison, self-report measures provided an accuracy of 76%. This mismatch may derive from measurement limitations for non-invasive neuroimaging, or perhaps from a fundamental lack of correspondence between heterogeneous clinical categories based on symptoms and neurobiological mechanisms.

## Methods

### Preregistration

The three main analyses in this manuscript were preregistered at https://osf.io/zndr2. Deviations are listed in **Supplementary Note 4.** The exploratory random forest analysis was added to test for possible nonlinear associations, due to evidence of its effectiveness in brain-behavior modelling,^41–43^ and finally, because rumination scores were non-normally distributed **(Extended Data** Fig. 1**, Supplementary** Fig. 4**, Supplementary Note 5)**. The external dataset was added in response to a reviewer request.

### Participants

The sample used in this study consisted of 486 English-speaking adolescents aged 12-18 recruited from the greater Boston area. After data quality screening (**Supplementary Note 6**), the final sample consisted of 444 adolescents (283 female, 161 male; Age *m* = 14.7 yrs, SD *=* 1.5 yrs). Participants were derived from four studies with similar baseline procedures, and one additional study added during review. Two of these studies (**Study 1, Study 2**) recruited adolescents who had elevated levels of anhedonia, as defined by an elevated anhedonia score (> 1) on the Kiddie Schedule for Affective Disorders and Schizophrenia (K-SADS^44^). Another study recruited adolescents with a parental history of MDD (**Study 3**). These three studies also recruited controls, with no lifetime psychiatric disorders (with the exception of ADHD in one study) or any use of psychotropic medication. In **Study 4,** ruminative adolescents were recruited, as defined by a score ≥ 13 on the Children’s Response Style Questionnaire^10^ (CRSQ) rumination subscale, with no control group. Finally, **Study 5** recruited adolescents with a parental history of MDD. Adolescents were ineligible from this study if they had a lifetime DSM-IV diagnosis of major depressive disorder, any anxiety disorder, or ADHD. Participants were ineligible for *all* studies if they had a lifetime [DSM-5] diagnosis of any psychotic disorder, bipolar disorder, past-year moderate or severe substance or alcohol use disorder, or chronic depression (episode > 2 years). Given the neuroimaging component of the studies, fMRI contradictions were also exclusionary. Additional exclusion criteria included hypothyroidism, medical or neurological illness (e.g., seizures), history of severe head injuries, and any developmental or intellectual disability that would interfere with a participant’s ability to complete study procedures. All psychotropic medications were exclusionary, except for stable doses of selective serotonin reuptake inhibitors (SSRIs) and serotonin and norepinephrine reuptake inhibitors (SNRIs). In summary, our sample consisted of 444 participants across five datasets, consisting of **Study 1** (n=79), **Study 2**(n=79), **Study 3** (n=116), **Study 4** (n=71) and **Study 5** (n=99). Full demographic details for each group are reported in **Supplementary Tables 1-5**. Pubertal development was assessed using the self-report Tanner Scale.^45^

All participants provided informed consent / assent and were compensated hourly for participation. All procedures were approved by the Mass General Brigham/ Partners IRB.

### Self-report Measures

The total score on the 13-item rumination subscale of the children’s response style questionnaire (CRSQ)^10^ was used to assess trait rumination. An example item is: “When I am sad, I think about how alone I feel.” Total scores reflected the sum of the items, and the internal reliability of the CRSQ was excellent (Cronbach’s alpha = .93). The external test dataset, **Study 5,** had considerably lower scores on the CRSQ (Welch’s *t*(225.36) = −7.62, *p <* 0.001, *d* = - 0.70). For the EMA measure of rumination, we operationalized rumination as ‘stress-reactive’, by inviting respondents to reflect on the most stressful experience in the last 24 hours or since the last EMA. Specifically, we used two questions “After this stressful or negative thing happened, I was dwelling on my mistakes, failures or losses”, “After this stressful or negative thing happened, I kept thinking about something negative that happened.” EMA responses were averaged over multiple days of sampling (**Supplementary Note 7**), For **Study 1 & 2,** EMA was reported on a 0-4 scale. For **Study 3 & 4**, EMA was reported on a 0-100 scale. To make the responses comparable, we z-scored them individually within each dataset, maintaining training and test independence^46^. For both measures, we removed outlier responses that were 2.5 SD above the mean, resulting distributions are summarized in **Supplementary Note 5**. EMA was not included in **Study 5**.

### fMRI Collection, Preprocessing, and Denoising

Resting-state imaging data was collected on 3T scanners. The overwhelming majority of scans from **Studies 1-4** were collected on the same PRISMA scanner, and **Study 5** was collected on a Tim Trio. For **Studies 1-4** two separate protocols were used consisting of 6-7 minute runs (see **Supplementary Note 8**). We analyzed each of the three runs from the **Study 4** protocol separately. **Study 5** involved single 5 minute scans (**Supplementary Note 8**). Preprocessing was conducted in SPM12 (http://fil.ion.ucl.ac.uk/spm/) and included grey matter segmentation, realignment and unwarping with a field map, slice-timing correction, co-registration, normalization to MNI space, resampling to 2 x 2 x 2 mm voxels, and smoothing with a 4 mm FWHM gaussian filter. Movement artifacts were screened and censored from subject-specific first-level models using Artifact Detection Tool (> 3 SD from mean intensity or > 1mm movement in any direction; http://www.nitrc.org/projects/artifact_detect/).

For denoising, we imported the SPM preprocessed data into the CONN toolbox (https://www.conn-toolbox.org/, v22a), after conducting an extra normalization step for the structural images and followed denoising steps from Kim et al. Data were band-pass filtered to include temporal frequencies between 0.008 Hz and 0.1 Hz. Nuisance covariates included 24 head motion parameters (six movement parameters including x, y, z, roll, pitch, and yaw, their mean-centered squares, their derivatives, and squared derivative), and the top five component scores from each of the white and cerebrospinal fluid (CSF) masks. We conducted despiking to reduce the impact of image-intensity outliers from ART (i.e., “spikes”). Any participants with more than 0.3 mm average framewise-displacement,^47^ were removed. This consisted of 6 subjects from **Study 1,** 1 subject from **Study 2,** 15 subjects from the largest **Study 3** dataset, 1 subject and 7 runs from **Study 4,** and 8 subjects from **Study 5.**

### fMRI Features

We extracted timecourses for all 443 participants from the 20 DMN seeds and all 280 Brainnetome regions (**Fig. 2**).^48^ We then calculated dynamic conditional correlations (DCC) between the timecourses using the *DCC_x_mex* function in the Dynamic Correlation toolbox (https://github.com/canlab/Lindquist_Dynamic_Correlation). We found that DCC was sensitive to scrubbing and scanner differences, and different DCC methods were differentially sensitive. We chose to implement DCC_x_mex and scrubbing, as this minimized the feature differences across the various datasets (**Supplementary** Fig. 5). The variance of DCC were our features, reflecting variability of interactions over time between brain regions. An example model (e.g. from the dMFPC seed) consisted of weights learned over 280 varDCC features (i.e., DMPFC to all other Brainnetome regions). We additionally examined mean DCC for the CPM approach (below). We averaged features within-subjects across clean runs in **Study 4**.

### Analysis Approaches

A schematic of the analyses approaches is shown in **Figures 1 and 3**. Our main analyses consisted of linear modelling approaches with feature selection, but we also created exploratory nonlinear random forest models. We examined predictive validity of our models in **Studies 1-4** using a training-validation split (all hyperparameter tuning was conducted in the training set). As only the nonlinear random forest models were significant in the validation dataset, for an external test of generalization we examined the performance of those models in **Study 5**. All modelling was conducted in MATLAB R2024a.

### Approach 1: Test DMPFC-Rum Model

The DMPFC-Rum Model consists of varDCC between the DMPFC and the rest of the brain, which predicts brooding in adults across multiple datasets (https://github.com/cocoanlab/rumination).^26^ Specifically, 84 weights were learned involving the inferior frontal gyrus, right frontal gyrus, cerebellum and more (across many brain networks) (**Fig. 3a**). Features were calculated in **Studies 1-4**, and then weights were applied to predict scores. We then correlated the predicted scores with the actual scores for both the CRSQ and EMA rumination measures, using Pearson’s correlations. We additionally examined the subgroups within the datasets. Finally, we tested whether predictions correlated with ratings of pubertal development to assess developmental invariance. The models were not assessed in **Study 5** as that dataset was set aside for testing generalizability of the random forest models.

### Approach 2: Lasso Models for all DMN Seeds

The approach herein is motivated by the DMPFC-Rum Model development,^26^ consisting of examining DCC between the 20 DMN seeds and 280 Brainnetome features (**Fig. 3b**). After calculating DCC features for the 20 seeds, we split **Studies 1-4** into an 80/20 train-test split, using the *caret* package in R. We balanced the splits on rumination scores, group, age, and sex. More detailed balancing was not achievable while maintaining an 80-20 ratio. However, as a post-hoc confirmation we tested the difference between puberty scores for each group and found no significant differences (*p* > .5).

To learn the mapping between the features and rumination, we conducted lasso regression with *alpha* parameter hyperoptimization, which controls how many weights are shrunk to zero. We evaluated 10-fold cross-validation within the internal dataset to calculate a correlation coefficient between predictions and actual values. We then shuffled the scores 10,000 times, conducted training again, and assessed the correlation coefficients. If the true correlation coefficient was higher than 95% of the permuted coefficients (*p* < .05), we judged that the model met our selection threshold. The selection threshold was not corrected for multiple comparisons. We then took selected models and tested them on the held-out data using Pearson’s correlations between predictions and test rumination scores. We considered null results in held-out data to be markers of a failure of the model to generalize.

### Approach 3: Whole-brain CPM

We considered whether regions besides the DMN could be linked to rumination (**Fig. 3c**). We used the same train-test splits from Approach 2, but now calculated pairwise dynamic connectivity for all 280 Brainnetome regions, resulting in 280*(280-1)/2 features. Connectome predictive modelling is a well-validated approach for whole-brain prediction,^39^ which consists of selecting correlated features and summing them into positive and negative networks. Models were selected based on a *p* < .05 threshold with permutation testing, and then selected models were tested in the test data. Controls for puberty and head motion were also assessed. Full details are present in **Supplementary Note 9**.

### Exploratory Models: Random Forest Regression

We explored whether nonlinear models (specifically, a decision-tree based random forest approach) provided more predictive power (**Fig. 3d**). We used the same train-test splits from Approaches 2 and 3. Models consisted of DMN-seeded models (20 models consisting of varDCC between each DMN seed and the 280 Brainnetome regions) and pairwise whole-brain models using mean connectivity and varDCC. Random forest models consisted of 250 trees with random partitions of features for each, with the number of features selected (*mtry)* as a hyperparameter using logarithmically spaced values from *mtry* =3 to *mtry* =148. Leaf size was set to a standard value of 5 (which controls the depth of the trees).^49^ All other parameters were set to Matlab defaults.

We calculated 10-fold cross-validation performance and then used a permutation shuffling procedure to select models, with 250 shuffles (the procedure was computationally intensive, preventing us from conducting more shuffles). If the true correlation coefficient was higher than 95% of the permuted coefficients (*p* < .05), we judged that the model met our selection threshold. We then took the selected models and tested the predictions using Pearson’s correlations in the held-out validation dataset, comparing the generalization performance to an uncorrected *p* < .05 threshold. No FDR correction was applied because of the subsequent independent test of generalizability.

Models meeting the threshold in the validation dataset were then additionally tested for generalizability using Pearson’s correlations in **Study 5.** Examination of the DCC features of Study 5 indicated widespread differences (e.g. mean levels) as well as differences for the most important features in the random forest models, due to differences in acquisition and scanner (**Supplementary Note 8**). Thus we applied COMBAT harmonization^50^ to the features, preserving differences related to rumination, sex, and puberty before testing the models. As this method is not often used with dynamic connectivity and may not fully retain nonlinear relationships with covariates, we also report results without the COMBAT harmonization in the Supplement. Finally, as Study 5 involved less ruminative adolescents, we examined sensitivity of the models using piecewise regression (for plotting) and piecewise correlations to assess if they performed better for rumination scores more similar to the training data.

For visualization only, we selected a random regression tree from the selected models to depict the branching rules. On the full models, we used out-of-bag importance maps to estimate feature contributions to the prediction performance. The maps were created using CANLab toolboxes (https://github.com/canlab/CanlabCore/). We compared feature importance across the 10 Brainnetome networks and compared the average importances to zero. Finally, we examined the role of participant factors such as puberty, age, sex, and clinical status in driving prediction performance.

## Data Availability

Data (feature values) will be available upon publication at https://osf.io/m5rgu/?view_only=b987fe5766f0488db763b3f70165a096.

## Code Availability

Code will be available upon publication at https://osf.io/m5rgu/?view_only=b987fe5766f0488db763b3f70165a096.

## Acknowledgements

This research was supported by NCCIH R01AT011002, NIMH R01MH116969, NIMH K23MH108752, the Tommy Fuss Fund, Klingenstein Third Generation Foundation, and a Young Investigator Grant from the Brain & Behavior Research Foundation awarded to CAW. Additionally, RPA received funding through the Tommy Fuss Fund, Dana Foundation, and Klingenstein Third Generation Foundation. We thank Paul Bloom, Carter Funkhouser and Julia Greenblatt for helpful suggestions and support. We thank Jungwoo Kim, Choong-Wan Woo and colleagues for making their model openly available for testing. We acknowledge the use of Claude 4.0 for code contributions for Supplemental Figures.

## Author Contributions

INT contributed conceptualization, formal analysis, investigation, methodology, project administration, software, visualization, writing-original draft and writing-editing. MP contributed investigation, visualization, writing-original draft and writing-editing. JS contributed formal analysis, investigation, and writing-editing. AT & NJ & KP contributed data curation and project administration. AK & JDG & RPA contributed writing-editing. CAW contributed funding acquisition, methodology, project administration, writing-editing, resources, and supervision.

## References

1. Mennin, D. S. & Fresco, D. M. What, me worry and ruminate about DSM 5 and RDoC? The importance of targeting negative self referential processing. Clin. Psychol. Sci. Pract. 20, 258–267 (2013).

2. Nolen-Hoeksema, S. Responses to depression and their effects on the duration of depressive episodes. J. Abnorm. Psychol. 100, 569–582 (1991).

3. Gong, T. et al. The associations among self criticism, hopelessness, rumination, and NSSI in adolescents: A moderated mediation model. J. Adolesc. 72, 1–9 (2019).

4. Mazzer, K., Boersma, K. & Linton, S. J. A longitudinal view of rumination, poor sleep and psychological distress in adolescents. J. Affect. Disord. 245, 686–696 (2019).

5. Surrence, K., Miranda, R., Marroquín, B. M. & Chan, S. Brooding and reflective rumination among suicide attempters: Cognitive vulnerability to suicidal ideation. Behav. Res. Ther. 47, 803–808 (2009).

6. McEvoy, P. M., Watson, H., Watkins, E. R. & Nathan, P. The relationship between worry, rumination, and comorbidity: Evidence for repetitive negative thinking as a transdiagnostic construct. J. Affect. Disord. 151, 313–320 (2013).

7. McLaughlin, K. A. & Nolen-Hoeksema, S. Rumination as a transdiagnostic factor in depression and anxiety. Behav. Res. Ther. 49, 186–193 (2011).

8. Hedegaard, H. & Warner, M. Suicide Mortality in the United States, 1999-2019. https://stacks.cdc.gov/view/cdc/101761(2021) doi:10.15620/cdc:101761.

9. Nolen-Hoeksema, S. & Morrow, J. A prospective study of depression and posttraumatic stress symptoms after a natural disaster: The 1989 Loma Prieta earthquake. J. Pers. Soc. Psychol. 61, 115–121 (1991).

10. Abela, J. R. Z., Brozina, K. & Haigh, E. P. An Examination of the Response Styles Theory of Depression in Third- and Seventh-Grade Children: A Short-Term Longitudinal Study. J. Abnorm. Child Psychol. 30, 515–527 (2002).

11. Abela, J. R. Z., Vanderbilt, E. & Rochon, A. A Test of the Integration of the Response Styles and Social Support Theories of Depression in Third and Seventh Grade Children. J. Soc. Clin. Psychol. 23, 653–674 (2004).

12. Moberly, N. J. & Watkins, E. R. Ruminative self-focus and negative affect: An experience sampling study. J. Abnorm. Psychol. 117, 314–323 (2008).

13. Ruscio, A. M. et al. Rumination predicts heightened responding to stressful life events in major depressive disorder and generalized anxiety disorder. J. Abnorm. Psychol. 124, 17–26 (2015).

14. Genet, J. J. & Siemer, M. Rumination moderates the effects of daily events on negative mood: Results from a diary study. Emotion 12, 1329–1339 (2012).

15. Zhou, H.-X. et al. Rumination and the default mode network: Meta-analysis of brain imaging studies and implications for depression. NeuroImage 206, 116287 (2020).

16. Raichle, M. E. et al. A default mode of brain function. Proc. Natl. Acad. Sci. 98, 676–682 (2001).

17. Kucyi, A., Kam, J. W. Y., Andrews-Hanna, J. R., Christoff, K. & Whitfield-Gabrieli, S. Recent advances in the neuroscience of spontaneous and off-task thought: implications for mental health. Nat. Ment. Health 1, 827–840 (2023).

18. Uddin, L. Q., Clare Kelly, A. M., Biswal, B. B., Xavier Castellanos, F. & Milham, M. P. Functional connectivity of default mode network components: Correlation, anticorrelation, and causality. Hum. Brain Mapp. 30, 625–637 (2009).

19. Fox, K. C. R., Spreng, R. N., Ellamil, M., Andrews-Hanna, J. R. & Christoff, K. The wandering brain: Meta-analysis of functional neuroimaging studies of mind-wandering and related spontaneous thought processes. NeuroImage 111, 611–621 (2015).

20. Ino, T., Nakai, R., Azuma, T., Kimura, T. & Fukuyama, H. Brain Activation During Autobiographical Memory Retrieval with Special Reference to Default Mode Network. Open Neuroimaging J. 5, 14–23 (2011).

21. Hamilton, J. P., Farmer, M., Fogelman, P. & Gotlib, I. H. Depressive Rumination, the Default-Mode Network, and the Dark Matter of Clinical Neuroscience. Biol. Psychiatry 78, 224–230 (2015).

22. Cooney, R. E., Joormann, J., Eugène, F., Dennis, E. L. & Gotlib, I. H. Neural correlates of rumination in depression. Cogn. Affect. Behav. Neurosci. 10, 470–478 (2010).

23. Berman, M. G. et al. Does resting-state connectivity reflect depressive rumination? A tale of two analyses. NeuroImage 103, 267–279 (2014).

24. Chen, X. et al. The subsystem mechanism of default mode network underlying rumination: A reproducible neuroimaging study. NeuroImage 221, 117185 (2020).

25. Tozzi, L. et al. Reduced functional connectivity of default mode network subsystems in depression: Meta-analytic evidence and relationship with trait rumination. NeuroImage Clin. 30, 102570 (2021).

26. Kim, J. et al. A dorsomedial prefrontal cortex-based dynamic functional connectivity model of rumination. Nat. Commun. 14, 3540 (2023).

27. Fettes, P. W. et al. Predictors and correlates of outcome for dorsolateral, dorsomedial, and orbitofrontal rTMS in major depression. Preprint at 10.21203/rs.3.rs-4484095/v1 (2024).

28. Marusak, H. A. et al. Mindfulness and dynamic functional neural connectivity in children and adolescents. Behav. Brain Res. 336, 211–218 (2018).

29. Treves, I. N. et al. Dynamic functional connectivity correlates of trait mindfulness in early adolescence. Biol. Psychiatry Glob. Open Sci. 100367 (2024) doi:10.1016/j.bpsgos.2024.100367.

30. Liégeois, R. et al. Resting brain dynamics at different timescales capture distinct aspects of human behavior. Nat. Commun. 10, 2317 (2019).

31. Chen (陈骁), X. & Yan (严超赣), C.-G. Hypostability in the default mode network and hyperstability in the frontoparietal control network of dynamic functional architecture during rumination. NeuroImage 241, 118427 (2021).

32. Hamilton, J. P. et al. Default-Mode and Task-Positive Network Activity in Major Depressive Disorder: Implications for Adaptive and Maladaptive Rumination. Biol. Psychiatry 70, 327– 333 (2011).

33. Kaiser, R. H. et al. Abnormal frontoinsular-default network dynamics in adolescent depression and rumination: a preliminary resting-state co-activation pattern analysis. Neuropsychopharmacology 44, 1604–1612 (2019).

34. Menon, V. Large-scale brain networks and psychopathology: a unifying triple network model. Trends Cogn. Sci. 15, 483–506 (2011).

35. Connolly, C. G. et al. Resting-State Functional Connectivity of Subgenual Anterior Cingulate Cortex in Depressed Adolescents. Biol. Psychiatry 74, 898–907 (2013).

36. Marchitelli, R. et al. Coupled changes between ruminating thoughts and resting-state brain networks during the transition into adulthood. Mol. Psychiatry 29, 3769–3778 (2024).

37. Gell, M., Noble, S., Laumann, T. O., Nelson, S. M. & Tervo-Clemmens, B. Psychiatric neuroimaging designs for individualised, cohort, and population studies. Neuropsychopharmacology (2024) doi:10.1038/s41386-024-01918-y.

38. Pujol, J. et al. Differences between the child and adult brain in the local functional structure of the cerebral cortex. NeuroImage 237, 118150 (2021).

39. Shen, X. et al. Using connectome-based predictive modeling to predict individual behavior from brain connectivity. Nat. Protoc. 12, 506–518 (2017).

40. Poldrack, R. A., Huckins, G. & Varoquaux, G. Establishment of Best Practices for Evidence for Prediction: A Review. JAMA Psychiatry 77, 534 (2020).

41. Agastinose Ronicko, J. F., et al. Diagnostic classification of autism using resting-state fMRI data improves with full correlation functional brain connectivity compared to partial correlation. J. Neurosci. Methods 345, 108884 (2020).

42. Kesler, S. R. et al. Predicting Long-Term Cognitive Outcome Following Breast Cancer with Pre-Treatment Resting State fMRI and Random Forest Machine Learning. Front. Hum. Neurosci. 11, 555 (2017).

43. Zhu, X., Du, X., Kerich, M., Lohoff, F. W. & Momenan, R. Random forest based classification of alcohol dependence patients and healthy controls using resting state MRI. Neurosci. Lett. 676, 27–33 (2018).

44. Kaufman, J. et al. Schedule for Affective Disorders and Schizophrenia for School-Age Children-Present and Lifetime Version (K-SADS-PL): Initial Reliability and Validity Data. J. Am. Acad. Child Adolesc. Psychiatry 36, 980–988 (1997).

45. Tanner, J. M. Growth at adolescence. Blackwell Sci. Publ. 2nd Edition, (1962).

46. Scheinost, D. et al. Ten simple rules for predictive modeling of individual differences in neuroimaging. NeuroImage 193, 35–45 (2019).

47. Power, J. D., Barnes, K. A., Snyder, A. Z., Schlaggar, B. L. & Petersen, S. E. Spurious but systematic correlations in functional connectivity MRI networks arise from subject motion. NeuroImage 59, 2142–2154 (2012).

48. Fan, L. et al. The Human Brainnetome Atlas: A New Brain Atlas Based on Connectional Architecture. Cereb. Cortex 26, 3508–3526 (2016).

49. Probst, P., Wright, M. N. & Boulesteix, A. Hyperparameters and tuning strategies for random forest. WIREs Data Min. Knowl. Discov. 9, e1301 (2019).

50. Fortin, J.-P. et al. Harmonization of cortical thickness measurements across scanners and sites. NeuroImage 167, 104–120 (2018).

51. Marek, S. et al. Reproducible brain-wide association studies require thousands of individuals. Nature 603, 654–660 (2022).

52. Spisak, T., Bingel, U. & Wager, T. D. Multivariate BWAS can be replicable with moderate sample sizes. Nature 615, E4–E7 (2023).

53. Tervo-Clemmens, B. et al. Reply to: Multivariate BWAS can be replicable with moderate sample sizes. Nature 615, E8–E12 (2023).

54. Hollon, S. D. What we got wrong about depression and its treatment. Behav. Res. Ther. 180, 104599 (2024).

55. Monroe, S. M. & Harkness, K. L. Major Depression and Its Recurrences: Life Course Matters. Annu. Rev. Clin. Psychol. 18, 329–357 (2022).

56. Fu, Z., Sui, J., Iraji, A., Liu, J. & Calhoun, V. Cognitive and Psychiatric Relevance of Dynamic Functional Connectivity States in a Large (N<10,000) Children Population. Preprint at 10.21203/rs.3.rs-3586731/v1 (2024).

57. Doucet, G. E. et al. Dev-Atlas: A reference atlas of functional brain networks for typically developing adolescents. Dev. Cogn. Neurosci. 72, 101523 (2025).

58. Kay, B. P. et al. Motion Impact Score for Detecting Spurious Brain-Behavior Associations. 2022.12.16.520797 Preprint at 10.1101/2022.12.16.520797 (2023).

59. Gratton, C. et al. Defining Individual-Specific Functional Neuroanatomy for Precision Psychiatry. Biol. Psychiatry 88, 28–39 (2020).

60. Khosla, M., Jamison, K., Ngo, G. H., Kuceyeski, A. & Sabuncu, M. R. Machine learning in resting-state fMRI analysis. Magn. Reson. Imaging 64, 101–121 (2019).

61. Taxali, A., Angstadt, M., Rutherford, S. & Sripada, C. Boost in Test–Retest Reliability in Resting State fMRI with Predictive Modeling. Cereb. Cortex 31, 2822–2833 (2021).

62. Dadi, K. et al. Benchmarking functional connectome-based predictive models for resting-state fMRI. NeuroImage 192, 115–134 (2019).

63. Herlin, B., Navarro, V. & Dupont, S. The temporal pole: From anatomy to function—A literature appraisal. J. Chem. Neuroanat. 113, 101925 (2021).

64. Olson, I. R., Plotzker, A. & Ezzyat, Y. The Enigmatic temporal pole: a review of findings on social and emotional processing. Brain 130, 1718–1731 (2007).

65. Olson, I. R., McCoy, D., Klobusicky, E. & Ross, L. A. Social cognition and the anterior temporal lobes: a review and theoretical framework. Soc. Cogn. Affect. Neurosci. 8, 123–133 (2013).

66. Kaiser, R. H., Andrews-Hanna, J. R., Wager, T. D. & Pizzagalli, D. A. Large-Scale Network Dysfunction in Major Depressive Disorder: A Meta-analysis of Resting-State Functional Connectivity. JAMA Psychiatry 72, 603 (2015).

67. Tse, N. Y. et al. A mega-analysis of functional connectivity and network abnormalities in youth depression. Nat. Ment. Health 2, 1169–1182 (2024).

68. He, P., Shi, Z., Cui, Y., Wang, R. & Wu, D. A spatiotemporal graph transformer approach for Alzheimer’s disease diagnosis with rs-fMRI. Comput. Biol. Med. 178, 108762 (2024).

69. Treves, I. N. et al. Connectome Based Predictive Modeling of Trait Mindfulness. Hum. Brain Mapp. 46, e70123 (2025).

70. Winter, N. R. et al. A Systematic Evaluation of Machine Learning–Based Biomarkers for Major Depressive Disorder. JAMA Psychiatry (2024) doi:10.1001/jamapsychiatry.2023.5083.

