## Supplementary Information for "Limited generalizability of dynamic fMRI correlates of adolescent rumination"

### Table of Contents

*Supplementary Text* ... *2*

**Supplementary Note 1. Factors Influencing Rumination and Related Brain Circuitry** ... 2

**Supplementary Note 2. Network Properties of Temporal Poles** ... 2

**Supplementary Note 3. External Validity Analyses** ... 2

**Supplementary Note 4. Deviations from Preregistration** ... 3

**Supplementary Note 5. Rumination Distributions** ... 3

**Supplementary Note 6. fMRI Quality Assurance Check** ... 4

**Supplementary Note 7. Ecological Momentary Assessment (EMA) Sampling** ... 4

**Supplementary Note 8. Scanner Protocols** ... 4

*Study 1-3 Collection* ... 4
*Study 4 Collection* ... 4
*Study 5 Collection* ... 5

**Supplementary Note 9. Connectome Predictive Modelling** ... 5

*Supplementary Tables* ... *6*

**Supplementary Table 1. Demographic and Clinical Characteristics of the Study 1 Sample** ... 6

**Supplementary Table 2. Demographic and Clinical Characteristics of the Study 2 Sample** ... 8

**Supplementary Table 3. Demographic and Clinical Characteristics of the Study 3 Sample** ... 10

**Supplementary Table 4. Demographic and Clinical Characteristics of the Study 4 Sample** ... 12

**Supplementary Table 5. Demographic and Clinical Characteristics of the Study 5 Sample** ... 14

**Supplementary Table 6. Correlations between model predictions and CRSQ rumination scores, by subgroup** ... 15

**Supplementary Table 7. Correlations between model predictions and EMA rumination scores, by subgroup** ... 16

**Supplementary Table 8. Full Lasso model reporting for CRSQ in training dataset** ... 17

**Supplementary Table 9. Full Lasso model reporting for EMA in training dataset** ... 18

**Supplementary Table 10. Non-parametric significance for CPM models in the training set** ... 19

**Supplementary Table 11. Non-parametric significance for random forest models predicting CRSQ in the training set** ... 20

**Supplementary Table 12: Network importances of Right Temporal Pole model** ... 21

**Supplementary Table 13. Non-parametric significance for random forest models predicting EMA in the training set** ... 22

**Supplementary Table 15. External test dataset performance for selected random forest models (with and without harmonization)** ... 24

*References* ... *33*

### Supplementary Figures

**Supplementary Figure 1. EMA Rumination score distribution across Studies 1-4** ... 25

**Supplementary Figure 2. Example Random Forest Decision Tree** ... 26

**Supplementary Figure 3. Whole-brain random forest model importances** ... 27

**Supplementary Figure 4. CRSQ Rumination score distribution in Study 5** ... 28

**Supplementary Figure 5. DMPFC seed dynamic conditional correlations** ... 29

**Supplementary Figure 6. Right temporal pole DCC differences between training and external test set for highest positive and negative importance regions** ... 30

**Supplementary Figure 7. Right temporal pole DCC differences between training and external test set for significant positive importance networks** ... 31

**Supplementary Figure 8. Static connectivity differences between training and external test set** ... 32

#### Supplementary Text

##### Supplementary Note 1. Factors Influencing Rumination and Related Brain Circuitry

Individual-level factors that affect rumination, depression, and related brain circuitry have been identified in adolescents. Sex differences emerge during this period, with girls showing a greater increase in brooding compared to boys,[1] which may account for the greater increase in depression in girls relative to boys during the adolescent years.[2] Previous research found a sex-specific association between salience network (SN) coherence and ruminative brooding in girls, where increased SN coherence, driven by dorsal anterior cingulate cortex (dACC) connectivity within the SN, was linked to higher levels of rumination.[3] Throughout puberty, significant changes occur in brain organization and network connectivity, which are closely tied to age and pubertal status.[4,5] This developmental period is characterized by heightened neural plasticity compared to adults, with pubertal youth exhibiting more dynamic changes in functional connectivity patterns as their brains mature. As a result, experiences during this time lead to more pronounced changes in neural circuitry.[6]

Parental genetic factors may contribute to the development of rumination and psychopathology in adolescents. Rumination has been shown to be moderately heritable and demonstrates shared genetic variability with depressive symptoms.[7] Environmental influences, such as negative life events, also play a role in shaping ruminative tendencies during adolescence,[8] which highlights the complex interplay between genetic and environmental factors in adolescent brain development. Moreover, rumination is comorbid with other depressive symptoms such as anhedonia, and both have been found to be associated with disruptions in cortico-limbic circuitry.[9,10]

##### Supplementary Note 2. Network Properties of Temporal Poles

We examined static functional connectivity for all participants in **Studies 1-4** between the 20 DMN seeds and 280 Fan connectome regions and calculated network properties of the seeds to investigate whether there were special features of the temporal poles. First, the left and right temporal poles showed on average smaller degrees than other DMN seeds ($M_L$ = 13.64, $M_{Ave}$ = 29.59, $t(344) = -23.29$, $p < 0.001$) and ($M_R$ =20.52, $M_{Ave}$ = 29.59, $t(344) = -11.59$, $p < 0.001$). They showed the same pattern in terms of decreased betweenness. The right temporal pole showed higher clustering coefficients than average DMN seeds ($M_R$ = 0.64, $M_{Ave}$ = 0.57, $t(344)$ = 5.24, $p < 0.001$). The left temporal pole showed lower local efficiency than average DMN seeds ($M_L$ = 0.63, $M_{Ave}$ = 0.72, $t(344) = -4.33$, $p < 0.001$).These results support the interpretation that the temporal poles are less connected and hub-like than the other DMN seeds and that the right temporal pole is especially locally connected and segregated.

##### Supplementary Note 3. External Validity Analyses

###### *External validation*

The external test dataset was harmonized using COMBAT, and then used to evaluate the selected random forest models (Left Temporal Pole, Right Temporal Pole, whole-brain). None were significant when examining the full distribution of rumination scores ($p$s > 0.35,

**Supplementary Table 15**), and there were no interactions with puberty (*p*s > 0.2). However, the external test dataset, **Study 5,** had considerably lower scores on the CRSQ (Welch's *t(*225.36) = -7.62, $p < 0.001$, $d = -0.70$). To account for this, we restricted analyses to CRSQ scores > 20, which corresponds to approximately 1 SD below the mean in the training set (it is reasonable to assume that the model fit from the training dataset is best for scores within 1 SD of the mean). We used 20 based on observations of the breakpoint in the distribution instead of the exact SD value. Given this admittedly arbitrary cut-off, the model showed a positive relationship that was similar in magnitude to the validation dataset ($r = 0.26$, $p = 0.064$).

More suggestive evidence of the value of the right temporal model comes from the non-harmonized dataset, where we found a significant ***negative*** relationship for the Right Temporal Pole model (**Supplementary Table 15)**, again suggesting that there is valuable information in that particular model but that factors like scanner, scan protocol, and population affect predictive performance. Differences between the external dataset and internal dataset connectivity distributions were assessed. Particularly notable were dynamic differences for highest positive importance regions (**Supplementary Fig. 6)**, which showed an average Cohen's $d = -0.45$ in the external dataset. For the significant networks (**Supplementary Fig. 7**), the DAN and Cerebellum showed even larger dynamic differences across datasets (*ds*= 0.68, 1.34, respectively), but the DMN was similar. Finally, static connectivity distributions were significantly different within these networks (**Supplementary Fig. 8)**. These results support the harmonization procedures here and suggest that improvements may be made in future work to account for static connectivity differences as well.

**Supplementary Note 4. Deviations from Preregistration**

- We added two additional datasets. One study, **Study 4,** is ongoing and similar in procedures to **Studies 1-3,** and we included the 71 adolescents in the training and validation datasets. During review, we added a completed external dataset consisting of 119 adolescents (99 after quality control) with CRSQ data, see **Study 5.**
- EMA responses were on different scales for **Study 1 & 2 vs Study 3 & 4,** so we z-scored within each dataset (maintaining test and train independence).
- We conducted puberty control analyses and conducted stratified training-test splits.
- We conducted an additional exploratory nonlinear analysis using seed-based analysis (DMN seeds), and upon reviewer request, a whole-brain nonlinear analysis.
- We did not conduct elastic net regression as it was deemed similar enough to the main lasso approach that it would not provide novel predictive power.
- We conducted connectome predictive modelling using the Brainnetome whole-brain atlas, not the Shen atlas, for comparability across methods.

**Supplementary Note 5. Rumination Distributions**

CRSQ scores were non-normally distributed across **all studies** ($W = 0.96$, $p < 0.001$) (**Extended Data Fig. 1** and **Supplementary Fig. 4**). EMA rumination scores were non-normally distributed across **Studies 1-4** ($W = 0.91$, $p < 0.001$), and not collected in **Study 5** (**Supplementary Fig. 1**).

#### Supplementary Note 6. fMRI Quality Assurance Check

Quality assurance (QA) checks were performed using the CONN Toolbox on all studies, to ensure data reliability. The preprocessing parameters were consistent with those employed by Kim et al., maintaining methodological consistency. Key metrics such as motion parameters, spatial normalization, and QC-FC were examined, and QA plots were constructed and reviewed to exclude any low-quality scans that could impact subsequent analyses.

#### Supplementary Note 7. Ecological Momentary Assessment (EMA) Sampling

For the **Study 1 & 2** datasets, we collected 2-3 EMA reports per day over 5 days. For **Study 3**, we collected EMA 4 times per day over 30 days. Finally, for **Study 4**, we collected EMA 3 times per day over 3 days. The differing timeframes reflect the different constraints of the studies – we sought to get EMA data that was relatively pure and free from intervention effects. In both studies, participants were first asked: “Think about the most stressful or negative thing that happened since you completed the last survey (or if this is your first survey, then the last 24 hours). At the worst point, how stressed did you feel?” and then asked the two rumination questions:

(1) “After this stressful or negative thing happened, I was dwelling on my mistakes, failures or losses”,

(2)“After this stressful or negative thing happened, I kept thinking about something negative that happened.”

**Study 5** did not include EMA.

#### Supplementary Note 8. Scanner Protocols

##### *Study 1-3 Collection*

Images were collected on two separate scanners, one Siemens Tim Trio 3.0 Tesla MRI with a 32-channel coil (for a small subset of **Study 1 & 2**), and one Siemens PRISMA 3.0 Tesla MRI with a 64-channel coil (for remaining **Study 1 & 2** and all of **Study 3**). The protocol was the same across scanners and consisted of a T1-weighted scan, one 6.5-minute resting-state scan, and a single gradient-echo fieldmap acquisition. Functional images were acquired with a multiband sequence (TR = 720 ms, TE = 30 ms, FOV = 212 mm, multiband accelerator factor = 6, voxel size = 2.5 × 2.5 × 2.5).

##### *Study 4 Collection*

Images were collected using a Siemens Prisma 3.0 Tesla MRI equipped with a 64-channel coil. The protocol consisted of a T1-weighted scan, three 6-minute, eyes-open, resting-state scans, and a single gradient-echo fieldmap acquisition. Functional images were acquired with a multiband sequence (TR = 720 ms, TE = 30 ms, FOV = 212 mm, multiband accelerator factor = 6, voxel size = 2.5 × 2.5 × 2.5).

##### *Study 5 Collection*

Images were collected using a Siemens Tim Trio 3.0 Tesla MRI equipped with a 32-channel coil. The protocol consisted of a T1-weighted scan, one five-minute eyes open resting-state scan, and a single gradient-echo fieldmap acquisition. Functional images were acquired with a multiband sequence (TR = 1300 ms, TE = 32.2 ms, FOV = 212 mm, multiband accelerator factor = 8, voxel size = 2.0 × 2.0 × 2.0).

#### Supplementary Note 9. Connectome Predictive Modelling

This procedure is similar to that followed by previous CPM studies.[11,12] We calculated correlations between CRSQ and EMA and the whole-brain features. We then selected edges with correlations that were below $p = 0.01$, assembling them into positive and negative networks, respectively. The positive network is summed to calculate "positive network strength" and the negative network is summed to calculate "negative network strength," and they may be subtracted to calculate "overall strength." They then may be correlated with rumination scores in held-out data. This was conducted 10-times (10-fold CV), and the correlation coefficient was compared to shuffled data. We shuffled the data 1000 times, and if the true correlation was greater than 95% of the shuffles ($p < 0.05$), we considered it to meet the selection threshold. For the selected models, we generated the positive and negative networks in a single fold and then tested these models in the held-out data. This procedure was also conducted using partial correlations. In analysis 1, we controlled for head motion. In analysis 2, we controlled for head motion and puberty. A final analysis consisted of conducting the same methods but using average DCC features, which are similar to static connectivity.[13]

#### Supplementary Tables

##### Supplementary Table 1. Demographic and Clinical Characteristics of the Study 1 Sample

| Sample Characteristics | S1 Anh (*n* = 38) | | S1 HC (*n* = 41) | | All S1 (*n* = 79) | |
|---|---|---|---|---|---|---|
| | **N** | **%** | **N** | **%** | **N** | **%** |
| **Biological Sex** | | | | | | |
| Female | 24 | 63.2 | 27 | 65.9 | 51 | 64.6 |
| Male | 14 | 36.8 | 14 | 34.1 | 28 | 35.4 |
| **Race** | | | | | | |
| American Indian | 0 | 0 | 0 | 0 | 0 | 0 |
| Asian | 3 | 7.9 | 4 | 9.8 | 7 | 8.9 |
| Black | 1 | 2.6 | 6 | 14.6 | 7 | 8.9 |
| Pacific | 0 | 0 | 1 | 2.4 | 1 | 1.3 |
| White | 30 | 78.9 | 28 | 68.3 | 58 | 73.4 |
| Multiracial | 4 | 10.5 | 2 | 4.9 | 6 | 7.6 |
| **Ethnicity** | | | | | | |
| Hispanic | 1 | 2.6 | 2 | 4.9 | 3 | 3.8 |
| Not Hispanic | 37 | 97.4 | 38 | 92.7 | 75 | 94.9 |
| Unknown | 0 | 0 | 1 | 2.4 | 1 | 1.3 |
| **Current Diagnoses (DSM-V)** | | | | | | |
| MDD | 23 | 60.5 | 0 | 0 | 23 | 29.1 |
| GAD | 9 | 23.7 | 0 | 0 | 9 | 11.4 |
| SAD | 5 | 13.2 | 0 | 0 | 5 | 6.3 |
| Panic Disorder | 1 | 2.6 | 0 | 0 | 1 | 1.3 |
| Specific Phobia | 1 | 2.6 | 0 | 0 | 1 | 1.3 |
| ADHD | 3 | 7.9 | 0 | 0 | 3 | 3.8 |
| ODD | 2 | 5.3 | 0 | 0 | 2 | 2.5 |
| PTSD | 1 | 2.6 | 0 | 0 | 1 | 1.3 |
| **Medication** | | | | | | |
| Psychotropic Medication | 7 | 18.4 | 0 | 0 | 7 | 8.9 |
| | **M (Range)** | **SD** | **M (Range)** | **SD** | **M (Range)** | **SD** |
| **Age (years)** | 15.82 (13 – 18) | 1.90 | 16.24 (13 – 18) | 1.70 | 16.04 (13 – 18) | 1.80 |
| **Family Income (dollars)** | 129,718.8 (20,000 – 350,000) | 67,317.88 | 151,685.8 (28,000 – 400,000) | 88,856.71 | 141,194.1 (20,000 – 400,000) | 79,486.67 |

*Note.* Anh: anhedonic individuals, selected by elevated scores on the KSADS interview. HC: healthy controls. MDD: Major Depressive Disorder. GAD: Generalized Anxiety Disorder. SAD:

Social Anxiety Disorder. ADHD: Attention-Deficit/Hyperactivity Disorder. ODD: Oppositional Defiant Disorder. PTSD: Post-Traumatic Stress Disorder.

**Supplementary Table 2. Demographic and Clinical Characteristics of the Study 2 Sample**

| Sample Characteristics | | | | | | |
|---|---|---|---|---|---|---|
| | **S2 Anh (*n* = 36)** | | **S2 HC (*n* = 43)** | | **All S2 (*n* = 79)** | |
| | **N** | **%** | **N** | **%** | **N** | **%** |
| **Biological Sex** | | | | | | |
| Female | 27 | 75.0 | 29 | 67.4 | 56 | 70.9 |
| Male | 9 | 25.0 | 14 | 32.6 | 23 | 29.1 |
| **Race** | | | | | | |
| American Indian | 0 | 0 | 0 | 0 | 0 | 0 |
| Asian | 4 | 11.1 | 6 | 14.0 | 10 | 12.7 |
| Black | 5 | 13.9 | 1 | 2.3 | 6 | 7.6 |
| Pacific | 1 | 2.8 | 0 | 0 | 1 | 1.3 |
| White | 19 | 52.8 | 30 | 69.8 | 49 | 62.0 |
| Multiracial | 4 | 11.1 | 5 | 11.6 | 9 | 11.4 |
| Other race | 2 | 5.6 | 1 | 2.3 | 3 | 3.8 |
| Unknown | 1 | 2.8 | 0 | 0 | 1 | 1.3 |
| **Ethnicity** | | | | | | |
| Hispanic | 6 | 16.7 | 4 | 9.3 | 10 | 12.7 |
| Not Hispanic | 30 | 83.3 | 39 | 90.7 | 69 | 87.3 |
| Unknown | 0 | 0 | 0 | 0 | 0 | 0 |
| **Current Diagnoses (DSM-V)** | | | | | | |
| MDD | 0 | 0 | 0 | 0 | 0 | 0 |
| GAD | 5 | 13.9 | 0 | 0 | 5 | 6.3 |
| SAD | 3 | 8.3 | 0 | 0 | 3 | 3.8 |
| Panic Disorder | 1 | 2.8 | 0 | 0 | 1 | 1.3 |
| Specific Phobia | 0 | 0 | 0 | 0 | 0 | 0 |
| ADHD | 1 | 2.8 | 1 | 2.3 | 2 | 2.5 |
| OCD | 2 | 5.6 | 0 | 0 | 2 | 2.5 |
| PTSD | 0 | 0 | 0 | 0 | 0 | 0 |
| **Medication** | | | | | | |
| Psychotropic Medication | 2 | 18.4 | 0 | 0 | 2 | 2.5 |
| | **M (Range)** | **SD** | **M (Range)** | **SD** | **M (Range)** | **SD** |
| **Age (years)** | 16.03 (12 – 18) | 2.09 | 15.84 (12 – 18) | 2.13 | 15.92 (12 – 18) | 2.10 |
| **Family Income (dollars)** | 146,424.20 (0 – 500,000) | 100,959.50 | 179,459.50 (40,000 – 400,000) | 81,817.75 | 163,885.70 (0 – 500,000) | 92,171.48 |

*Note.* Anh: anhedonic individuals, selected by elevated scores on the KSADS interview. HC: healthy controls. MDD: Major Depressive Disorder. GAD: Generalized Anxiety Disorder. SAD: Social Anxiety Disorder. ADHD: Attention-Deficit/Hyperactivity Disorder. OCD: Obsessive Compulsive Disorder. PTSD: Post-Traumatic Stress Disorder.

**Supplementary Table 3. Demographic and Clinical Characteristics of the Study 3 Sample**

| **Sample Characteristics** | | | | | | |
|---|---|---|---|---|---|---|
| | **S3 HR (*n* = 49)** | | **S3 LR (*n* = 67)** | | **All S3 (*n* = 116)** | |
| | **N** | **%** | **N** | **%** | **N** | **%** |
| **Biological Sex** | | | | | | |
| Female | 31 | 63.3 | 28 | 41.8 | 59 | 50.9 |
| Male | 18 | 36.7 | 39 | 58.2 | 57 | 49.1 |
| **Race** | | | | | | |
| American Indian | 1 | 2.04 | 0 | 0 | 1 | 0.86 |
| Asian | 0 | 0 | 3 | 4.48 | 3 | 2.59 |
| Black | 1 | 2.04 | 2 | 2.99 | 3 | 2.59 |
| Pacific | 0 | 0 | 0 | 0 | 0 | 0 |
| White | 41 | 83.7 | 55 | 82.1 | 96 | 82.8 |
| Multiracial | 5 | 10.2 | 7 | 10.4 | 12 | 10.3 |
| Unknown | 1 | 2.04 | 0 | 0 | 1 | 0.86 |
| **Ethnicity** | | | | | | |
| Hispanic | 7 | 14.3 | 2 | 2.99 | 9 | 7.76 |
| Not Hispanic | 42 | 85.7 | 65 | 97.0 | 107 | 92.2 |
| **Current Diagnoses (DSM-V)** | | | | | | |
| GAD | 1 | 2.04 | 0 | 0 | 1 | 0.86 |
| Specific Phobia | 1 | 2.04 | 0 | 0 | 1 | 0.86 |
| ODD | 0 | 0 | 0 | 0 | 0 | 0 |
| OCD | 1 | 2.04 | 0 | 0 | 1 | 0.86 |
| PTSD | 1 | 2.04 | 0 | 0 | 1 | 0.86 |
| **Medication** | | | | | | |
| Psychotropic Medication | 0 | 0 | 0 | 0 | 0 | 0 |
| | **M (Range)** | **SD** | **M (Range)** | **SD** | **M (Range)** | **SD** |
| **Age (years)** | 13.5 (12-15) | 1.02 | 13.6 (12-15) | 1.09 | 13.6 (12-15) | 1.1 |
| **Family Income (dollars)** | 163,532 (20,500-900,000) | 184,269.7 | 143,414 (45,000-650,000) | 110,582.0 | 151,078 (20,500-900,000) | 142,451.9 |

*Note.* HR: High-Risk. LR: Low-Risk. GAD: Generalized Anxiety Disorder. ODD: Oppositional Defiant Disorder. OCD: Obsessive Compulsive Disorder. PTSD: Post-Traumatic Stress Disorder. The following diagnoses were not included due to no participants having met criteria, Major Depressive Episode, Social Anxiety Disorder, Panic Disorder, Selective Mutism,

Attention-Deficit/Hyper-Activity Disorder, Binge Eating Disorder, Alcohol/Substance-Use Disorders, Oppositional Defiant Disorder.

**Supplementary Table 4. Demographic and Clinical Characteristics of the Study 4 Sample**

| Sample Characteristics | | |
|---|---|---|
| | **N (*n* = 71)** | **%** |
| **Biological Sex** | | |
| Female | 53 | 74.6 |
| Male | 18 | 25.4 |
| **Race** | | |
| American Indian | 1 | 1.4 |
| Asian | 9 | 12.7 |
| Black | 5 | 7.0 |
| Pacific | 1 | 1.4 |
| White | 43 | 60.6 |
| Multiracial | 9 | 12.7 |
| Unknown | 3 | 4.2 |
| **Ethnicity** | | |
| Hispanic or Latino | 9 | 12.7 |
| Not Hispanic or Latino | 62 | 87.3 |
| **Current Diagnoses (DSM-V)** | | |
| MDD | 6 | 8.5 |
| GAD | 31 | 43.7 |
| SAD | 20 | 28.2 |
| Panic Disorder | 7 | 9.9 |
| Specific Phobia | 5 | 7.0 |
| Selective Mutism | 1 | 1.4 |
| ADHD | 1 | 1.4 |
| ODD | 3 | 4.2 |
| OCD | 2 | 2.8 |
| Binge Eating Disorder | 3 | 4.2 |
| PTSD | 1 | 1.4 |
| Alcohol / Substance Use Disorders | 0 | 0.0 |
| **Medication** | | |
| Psychotropic Medication | 4 | 5.6 |
| **Handedness** | | |
| Right | 44 | 62.0 |
| Left | 7 | 9.9 |
| Unknown | 20 | 28.2 |
| | **M (Range)** | **SD** |
| **Age (years)** | 15.8 (13 -18) | 1.6 |

| | | |
|---|---|---|
| **Family Income (dollars)** | 186,017.9 (3,000 - 500,000) | 113,751.3 |

*Note.* One participant was excluded for head motion, but was included here for ease of reporting. MDD: Major Depressive Disorder. GAD: Generalized Anxiety Disorder. SAD: Social Anxiety Disorder. ADHD: Attention-Deficit/Hyperactivity Disorder. ODD: Oppositional Defiant Disorder. OCD: Obsessive Compulsive Disorder. PTSD: Post-Traumatic Stress Disorder.

**Supplementary Table 5. Demographic and Clinical Characteristics of the Study 5 Sample**

| Sample Characteristics | | |
|---|---|---|
| | **N (*n* = 99)** | **%** |
| **Biological Sex** | | |
| Female | 64 | 64.6 |
| Male | 35 | 35.4 |
| **Race** | | |
| American Indian | 0 | 0.0 |
| Asian | 5 | 5.1 |
| Black | 3 | 3.0 |
| Pacific | 0 | 0.0 |
| White | 81 | 81.8 |
| Multiracial | 10 | 10.1 |
| Unknown | 0 | 0.0 |
| **Ethnicity** | | |
| Hispanic or Latino | 5 | 5.1 |
| Not Hispanic or Latino | 94 | 94.9 |
| **Current Diagnoses (DSM-V)** | | |
| All | 0 | 0.0 |
| **Family Income (dollars)** | | |
| 50,000 – 75,000 | 8 | 8.1 |
| 75,000 – 100,000 | 16 | 16.2 |
| Greater than 100,000 | 65 | 65.7 |
| Unknown | 10 | 10.0 |
| | **M (Range)** | **SD** |
| **Age (years)** | 13.0 (12 -14) | 0.8 |

*Note*. All participants were right-handed. Due to exclusionary criteria, no participants were taking psychotropic medications, and no participants met threshold criteria for a current psychiatric diagnosis (DSM-IV). MDD: Major Depressive Disorder. GAD: Generalized Anxiety Disorder. SAD: Social Anxiety Disorder. ADHD: Attention-Deficit/Hyperactivity Disorder. ODD: Oppositional Defiant Disorder. OCD: Obsessive Compulsive Disorder. PTSD: Post-Traumatic Stress Disorder.

**Supplementary Table 6. Correlations between model predictions and CRSQ rumination scores, by subgroup**

| Group | Sample size, n (denoised) | R-value | P-value | FDR-p |
|---|---|---|---|---|
| S1_Anh | 37 | -.03 | .86 | .97 |
| S1_HC | 40 | .13 | .41 | .72 |
| S2_Anh | 28 | .01 | .97 | .97 |
| S2_HC | 40 | -.07 | .67 | .94 |
| S3_HR | 48 | .14 | .34 | .72 |
| S3_HC | 67 | -.13 | .31 | .72 |
| S4 | 69 | .20 | .10 | .69 |

S1-S4: samples 1 to 4. Anh: anhedonic based on KSADS score. HR: high risk, parental history of MDD. * $p < .05$.

**Supplementary Table 7. Correlations between model predictions and EMA rumination scores, by subgroup**

| Group | Sample size, n (denoised) | R-value | P-value | FDR-p |
|---|---|---|---|---|
| S1_Anh | 37 | -.23 | .16 | .30 |
| S1_HC | 41 | .02 | .88 | .91 |
| S2_Anh | 31 | -.25 | .17 | .30 |
| S2_HC | 38 | .16 | .35 | .49 |
| S3_HR | 48 | -.02 | .91 | .91 |
| S3_HC | 64 | -.28 | **.02*** | .16 |
| S4 | 68 | .25 | **.04*** | .16 |

S1-S4: samples 1 to 4. Anh: anhedonic based on KSADS score. HR: high risk, parental history of MDD. * $p < .05$.

**Supplementary Table 8. Full Lasso model reporting for CRSQ in training dataset**

| **Seed Area** | ***p*-values** | **FDR-*p*** |
|---|---|---|
| DMN_l_HF | 0.2232 | 0.576 |
| DMN_l_LTC | 0.1901 | 0.576 |
| DMN_l_PCC | 0.6066 | 0.6066 |
| DMN_l_PHC | 0.1025 | 0.576 |
| DMN_l_Rsp | 0.2543 | 0.576 |
| DMN_l_TPJ | 0.5425 | 0.6066 |
| DMN_l_TempP | 0.5183 | 0.6066 |
| DMN_l_aMPFC | 0.4171 | 0.6066 |
| DMN_l_pIPL | 0.5698 | 0.6066 |
| DMN_r_HF | 0.2098 | 0.576 |
| DMN_r_LTC | 0.3168 | 0.576 |
| DMN_r_PCC | 0.4123 | 0.6066 |
| DMN_r_PHC | 0.0314* | 0.576 |
| DMN_r_Rsp | 0.2047 | 0.576 |
| DMN_r_TPJ | 0.5955 | 0.6066 |
| DMN_r_TempP | 0.2981 | 0.576 |
| DMN_r_aMPFC | 0.3102 | 0.576 |
| DMN_r_pIPL | 0.5985 | 0.6066 |
| DMN_vMPFC | 0.0684 | 0.576 |
| DMN_DMPFC | 0.4529 | 0.6066 |

Non-parametric *ps* for lasso models trained on dynamic connectivity from the seeds to the Brainnetome regions to predict CRSQ. * $p < 0.05$, passes selection threshold

**Supplementary Table 9. Full Lasso model reporting for EMA in training dataset**

| **Area** | ***p*-values** | **FDR-p** |
|---|---|---|
| DMN_l_HF | 0.3588 | 0.650 |
| DMN_l_LTC | 0.2042 | 0.650 |
| DMN_l_PCC | 0.1619 | 0.650 |
| DMN_l_PHC | 0.366 | 0.650 |
| DMN_l_Rsp | 0.9245 | 0.994 |
| DMN_l_TPJ | 0.7521 | 0.956 |
| DMN_l_TempP | 0.9482 | 0.994 |
| DMN_l_aMPFC | 0.1855 | 0.650 |
| DMN_l_pIPL | 0.3902 | 0.650 |
| DMN_r_HF | 0.7801 | 0.956 |
| DMN_r_LTC | 0.8008 | 0.956 |
| DMN_r_PCC | 0.0109* | 0.218 |
| DMN_r_PHC | 0.8124 | 0.956 |
| DMN_r_Rsp | 0.3753 | 0.650 |
| DMN_r_TPJ | 0.1282 | 0.650 |
| DMN_r_TempP | 0.9944 | 0.994 |
| DMN_r_aMPFC | 0.3572 | 0.650 |
| DMN_r_pIPL | 0.7323 | 0.956 |
| DMN_vMPFC | 0.2566 | 0.650 |
| DMN_DMPFC | 0.3688 | 0.650 |

Non-parametric *ps* for lasso models trained on dynamic connectivity from the seeds to the Brainnetome regions to predict EMA. * $p < 0.05$, passes selection threshold

**Supplementary Table 10. Non-parametric significance for CPM models in the training set**

| Measure | Pos Network | Neg network | Both | MEAN Connectivity |
|---|---|---|---|---|
| *No partials* | | | | |
| CRSQ RUM | -0.13 | 0.11 (*p* = 0.033) * | 0.12 (*p* = 0.063) | -0.076 |
| EMA RUM | -0.0014 | 0.025 | 0.033 | 0.10 |
| *Partial HM* | | | | |
| CRSQ RUM | -0.13 | 0.12 (*p* = 0.025)* | 0.12 (*p* = 0.043)* | -0.054 |
| EMA RUM | -0.030 | 0.062 | 0.07 | 0.10 |
| *Partial HM and puberty* | | | | |
| CRSQ RUM | -0.17 | 0.18 (*p* = 0.004)** | 0.19 (*p* = 0.006)** | -0.021 |
| EMA RUM | -0.069 | 0.089 (p = 0.075) | 0.095 | 0.10 |

Models were trained on dynamic and mean connectivity from all Brainnetome regions to predict CRSQ and EMA. Mean connectivity column shows overall network performance (pos-neg). One positive network model for mean connectivity passed selection threshold ($p = 0.0375$), but did not generalize. Only models that passed selection threshold display p-values, * $p < 0.05$, ** $p < 0.01$. HM : head motion, puberty: total puberty score on tanner scale.

**Supplementary Table 11. Non-parametric significance for random forest models predicting CRSQ in the training set**

| **Area** | ***p*-value** | **FDR-*p*** |
|---|---|---|
| DMN_l_HF | 0.004* | 0.013 |
| DMN_l_LTC | 0.008* | 0.018 |
| DMN_l_PCC | 0.004* | 0.013 |
| DMN_l_PHC | 0.188 | 0.221 |
| DMN_l_Rsp | 0.564 | 0.564 |
| DMN_l_TPJ | 0.008* | 0.018 |
| DMN_l_TempP | 0.004* | 0.013 |
| DMN_l_aMPFC | 0.012 | 0.024 |
| DMN_l_pIPL | 0.084 | 0.129 |
| DMN_r_HF | 0.340 | 0.358 |
| DMN_r_LTC | 0.104 | 0.149 |
| DMN_r_PCC | 0.004* | 0.013 |
| DMN_r_PHC | 0.004* | 0.013 |
| DMN_r_Rsp | 0.284 | 0.316 |
| DMN_r_TPJ | 0.016* | 0.029 |
| DMN_r_TempP | 0.048* | 0.080 |
| DMN_r_aMPFC | 0.172 | 0.220 |
| DMN_r_pIPL | 0.008* | 0.018 |
| DMN_vMPFC | 0.004* | 0.013 |
| DMN_DMPFC | 0.176 | 0.220 |

* $p < 0.05$, passes selection threshold.

**Supplementary Table 12: Network importances of Right Temporal Pole model**

| Networks | Mean Importances | P-value | FDR-P |
|---|---|---|---|
| Brainstem | 0.019 | 0.678 | 0.678 |
| Cerebellum | 0.028 | 0.034* | 0.112 |
| dAttention | 0.031 | 0.014* | 0.068 |
| Default | 0.028 | 0.013* | 0.068 |
| Frontopariet al | 0.021 | 0.079 | 0.197 |
| Limbic | -0.011 | 0.348 | 0.435 |
| Somatomoto r | 0.020 | 0.149 | 0.266 |
| Subcortical | -0.015 | 0.218 | 0.311 |
| vAttention | 0.008 | 0.448 | 0.498 |
| Visual | 0.015 | 0.159 | 0.266 |

Importances for all regions in a network were statistically compared to zero with single sample *t*-tests , * $p < 0.05$

**Supplementary Table 13. Non-parametric significance for random forest models predicting EMA in the training set**

| **Area** | **p-values** | **FDR-p** |
|---|---|---|
| DMN_l_HF | 0.752 | 0.904 |
| DMN_l_LTC | 0.168 | 0.480 |
| DMN_l_PCC | 0.060 | 0.240 |
| DMN_l_PHC | 0.724 | 0.904 |
| DMN_l_Rsp | 0.500 | 0.855 |
| DMN_l_TPJ | 0.556 | 0.855 |
| DMN_l_TempP | 0.932 | 0.981 |
| DMN_l_aMPFC | 0.044* | 0.220 |
| DMN_l_pIPL | 0.132 | 0.440 |
| DMN_r_HF | 0.740 | 0.904 |
| DMN_r_LTC | 0.768 | 0.904 |
| DMN_r_PCC | 0.040* | 0.220 |
| DMN_r_PHC | 0.524 | 0.855 |
| DMN_r_Rsp | 0.400 | 0.855 |
| DMN_r_TPJ | 0.004* | 0.040 |
| DMN_r_TempP | 0.992 | 0.992 |
| DMN_r_aMPFC | 0.004* | 0.040 |
| DMN_r_pIPL | 0.920 | 0.981 |
| DMN_vMPFC | 0.216 | 0.540 |
| DMN_DMPFC | 0.552 | 0.855 |

* $p < 0.05$, passes selection threshold. Held-out data performance of selected models was non-significant.

**Supplementary Table 14. Non-parametric significance for whole-brain random forest models predicting CRSQ and EMA in training set.**

| Measure | Both | MEAN Connectivity |
|---|---|---|
| CRSQ RUM | 0.12 ($p$ = 0.012*) | 0.13 ($p$ = 0.012*) |
| EMA RUM | -0.02 ($p$ = 0.38) | 0.02 ($p$ = 0.16) |

* $p < 0.05$, passes selection threshold

**Supplementary Table 15. External test dataset performance for selected random forest models (with and without harmonization)**

| Model | Harmonized | Non-harmonized |
|---|---|---|
| LTP | 0.05 (*p* = 0.61) | -0.02 (*p* = 0.84) |
| RTP | 0.09 (*p* = 0.39) | -0.23* (*p* = 0.02) |
| Whole-brain | -0.01 (*p* = 0.85) | -0.04 (*p* = 0.67) |

Sample size = 99. Pearson's correlations and *p* values (parentheses) are shown. * $p < 0.05$

#### Supplementary Figures

##### Supplementary Figure 1. EMA Rumination score distribution across Studies 1-4

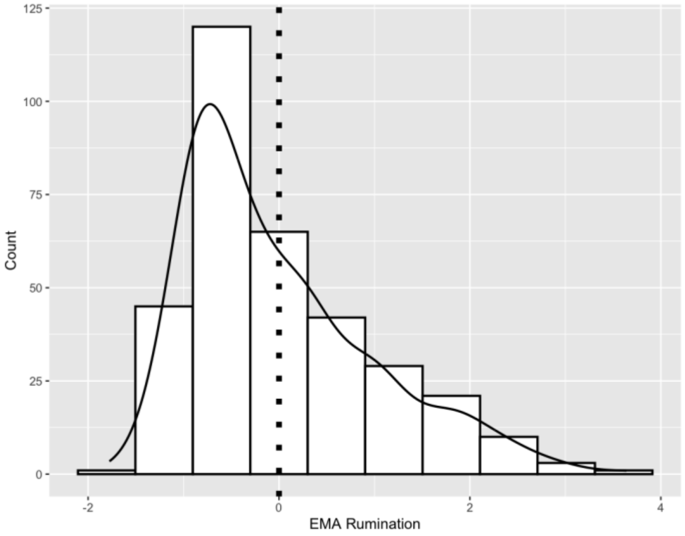


EMA: ecological probes of rumination, Count: number of participants. Dotted line represents mean (*n* = 344). Values are z-scored within datasets (maintaining train-test independence). No EMA was present in **Study 5.**

**Supplementary Figure 2. Example Random Forest Decision Tree**

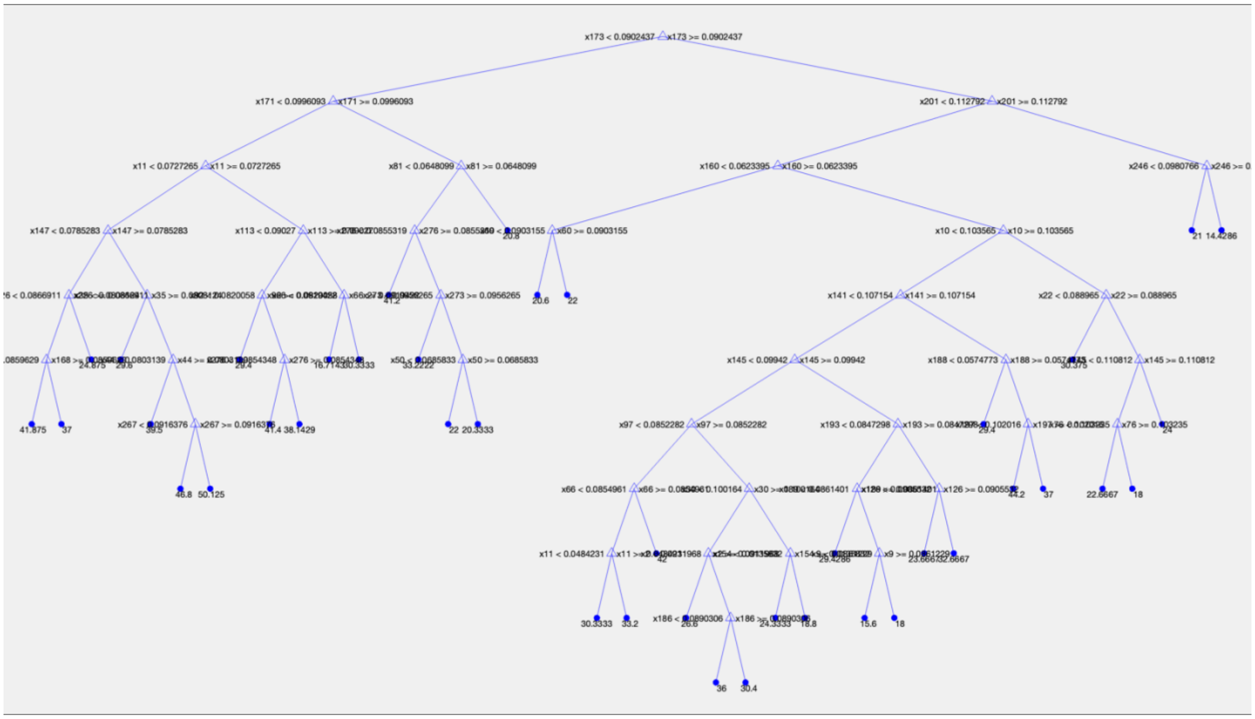


A regression tree from the left temporal pole model. Each branch reflects a rule based on varDCC for that feature (e.g. the value of varDCC for temporal pole and node 270). The final values are the outputted predictions.

**Supplementary Figure 3. Whole-brain random forest model importances**

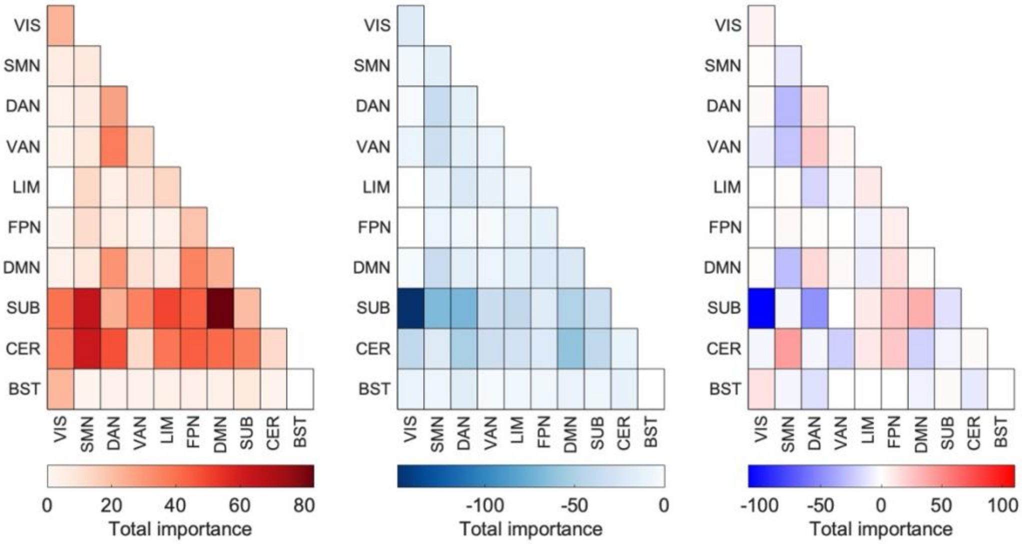


Total importance is the total weight of all out-of-bag importances from the random forest model for a given set of connections between networks from the Brainnetome atlas. On left, positive sum weights, middle negative sum weights, and right overall sum weights. More red represents higher positive weights, more blue represents lower negative weights.

**Supplementary Figure 4. CRSQ Rumination score distribution in Study 5**

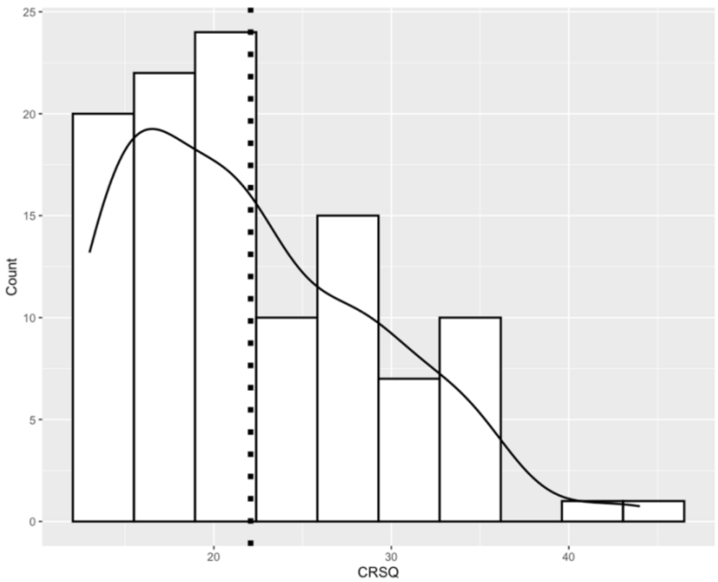


CRSQ: child response style questionnaire, Count: number of participants. Dotted line represents mean (*n* = 99 participants).

**Supplementary Figure 5. DMPFC seed dynamic conditional correlations**

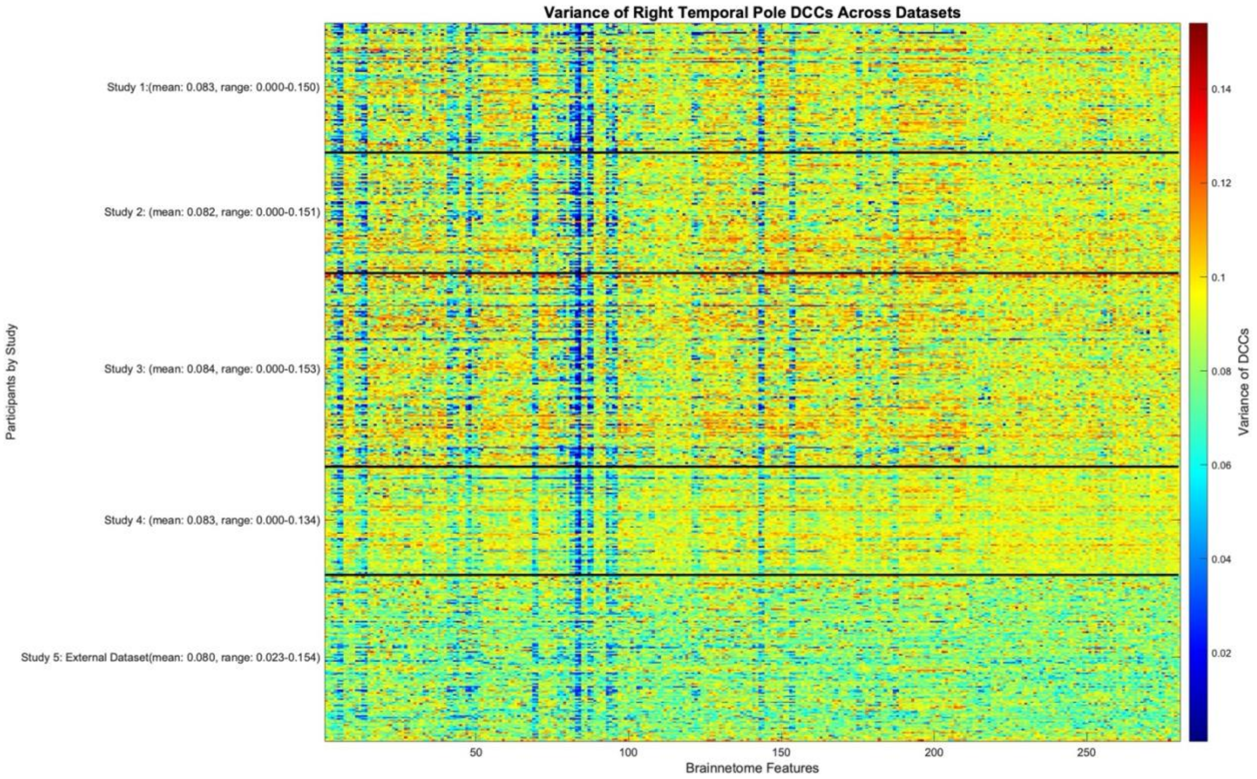


Example Right Temporal Pole dynamic connectivity features across the 443 subjects (y-axis) and 280 Brainnetome regions (x-axis). **Studies 4 and 5** involved different scan protocols. Red represents higher variance of dynamic connectivity, blue represents lower variance.

**Supplementary Figure 6. Right temporal pole DCC differences between training and external test set for highest positive and negative importance regions**

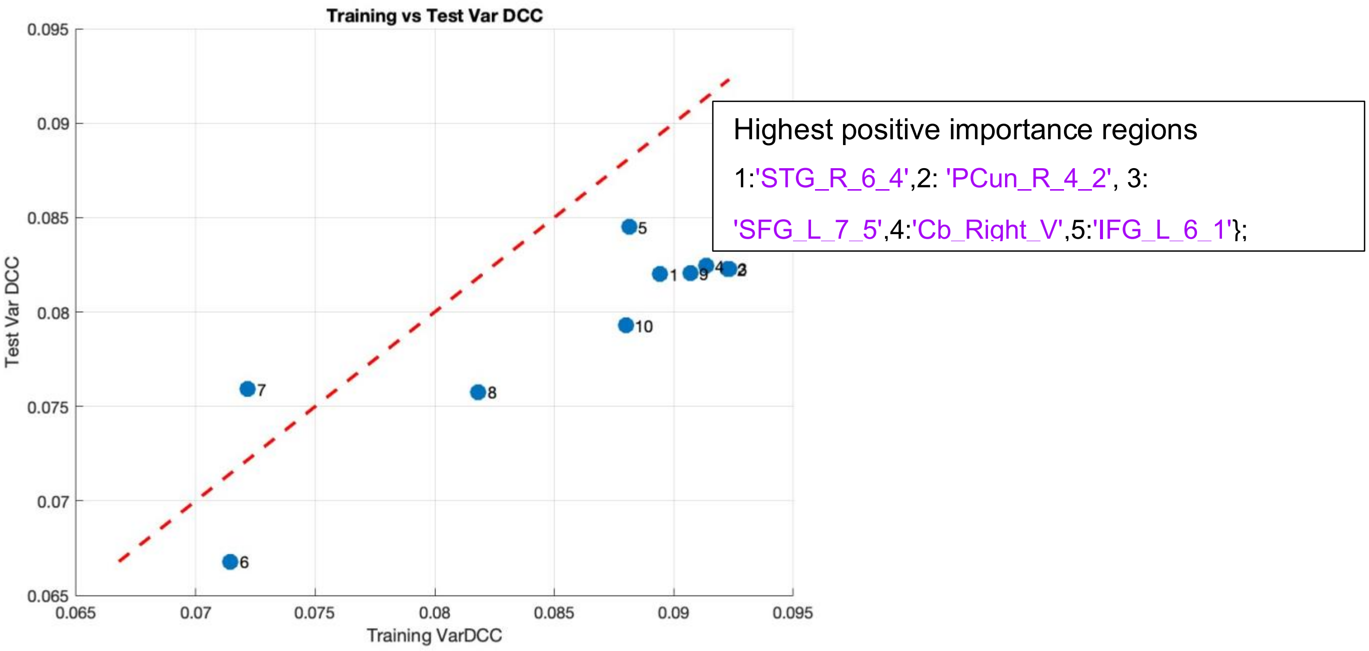


Points represent Training (x-axis) and External Test (y-axis) feature values for high importance regions. The red line represents equal feature values. Smaller DCC values may be observed for the regions in the external test set.

**Supplementary Figure 7. Right temporal pole DCC differences between training and external test set for significant positive importance networks**

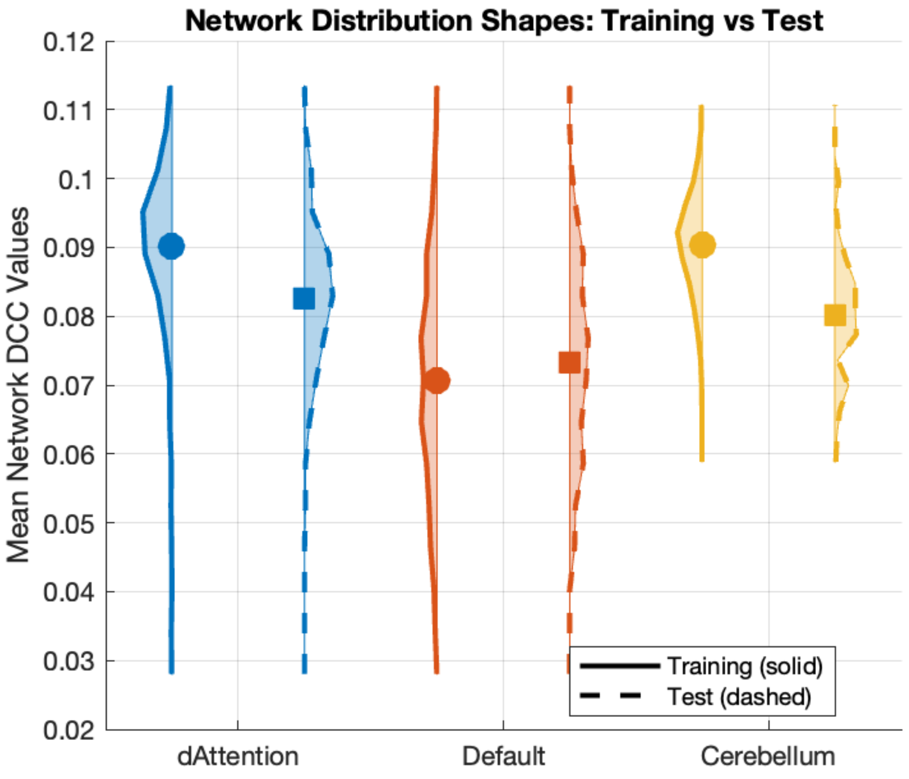


High importance networks shown for training and external data (test) splits. The variance of dynamic connectivity is averaged across all RTP – network edges.

**Supplementary Figure 8. Static connectivity differences between training and external test set**

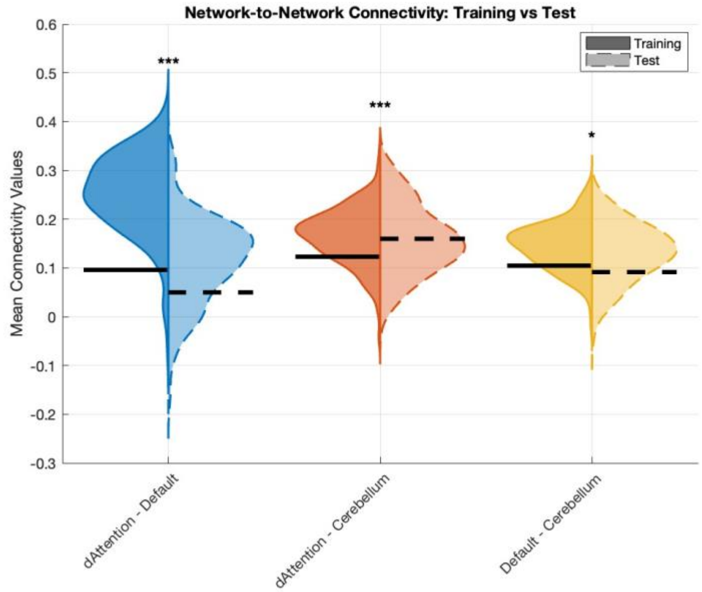


Network static connectivity differences for training and external test set. Connections between high importance networks are shown as they exemplify differences. $p < 0.05$ *** $p < 0.001$

**References**

1. Lopez, C. M., Driscoll, K. A. & Kistner, J. A. Sex Differences and Response Styles: Subtypes of Rumination and Associations with Depressive Symptoms. *J. Clin. Child Adolesc. Psychol.* **38**, 27–35 (2009).
2. Burwell, R. A. & Shirk, S. R. Subtypes of Rumination in Adolescence: Associations Between Brooding, Reflection, Depressive Symptoms, and Coping. *J. Clin. Child Adolesc. Psychol.* **36**, 56–65 (2007).
3. Ordaz, S. J. *et al.* Ruminative brooding is associated with salience network coherence in early pubertal youth. *Soc. Cogn. Affect. Neurosci.* **12**, 298–310 (2017).
4. Pelletier-Baldelli, A. *et al.* Brain network connectivity during peer evaluation in adolescent females: Associations with age, pubertal hormones, timing, and status. *Dev. Cogn. Neurosci.* **66**, 101357 (2024).
5. Tamnes, C. K. *et al.* Development of the Cerebral Cortex across Adolescence: A Multisample Study of Inter-Related Longitudinal Changes in Cortical Volume, Surface Area, and Thickness. *J. Neurosci.* **37**, 3402–3412 (2017).
6. Sydnor, V. J. & Satterthwaite, T. D. Neuroimaging of plasticity mechanisms in the human brain: from critical periods to psychiatric conditions. *Neuropsychopharmacology* **48**, 219–220 (2023).
7. Johnson, D. P., Whisman, M. A., Corley, R. P., Hewitt, J. K. & Friedman, N. P. Genetic and environmental influences on rumination and its covariation with depression. *Cogn. Emot.* **28**, 1270–1286 (2014).
8. Chen, J. & Li, X. Genetic and Environmental Influences on Adolescent Rumination and its Association with Depressive Symptoms. *J. Abnorm. Child Psychol.* **41**, 1289–1298 (2013).

9. Burkhouse, K. L. *et al.* Neural correlates of rumination in adolescents with remitted major depressive disorder and healthy controls. *Cogn. Affect. Behav. Neurosci.* **17**, 394–405 (2017).

10. Young, K., Sandman, C. & Craske, M. Positive and Negative Emotion Regulation in Adolescence: Links to Anxiety and Depression. *Brain Sci.* **9**, 76 (2019).

11. Kucyi, A. *et al.* Prediction of stimulus-independent and task-unrelated thought from functional brain networks. *Nat. Commun.* **12**, 1793 (2021).

12. Treves, I. N. *et al.* Connectome predictive modeling of trait mindfulness. 2024.07.09.602725 Preprint at https://doi.org/10.1101/2024.07.09.602725 (2024).

13. Kim, J. *et al.* A dorsomedial prefrontal cortex-based dynamic functional connectivity model of rumination. *Nat. Commun.* **14**, 3540 (2023).
