## Extended Data for "Limited generalizability of dynamic fMRI correlates of adolescent rumination"

Extended Data Figures for Preregistration

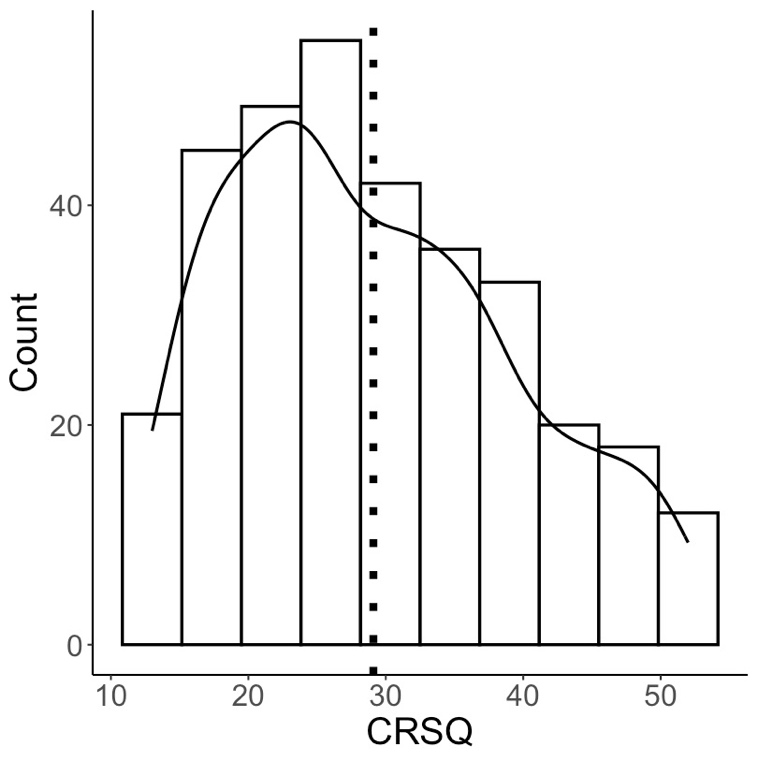

Figure 1: Rumination distributions for internal data (Studies 1 to 4). CRSQ: Child Response Style Questionnaire, Count: number of participants. Dotted line represents mean.

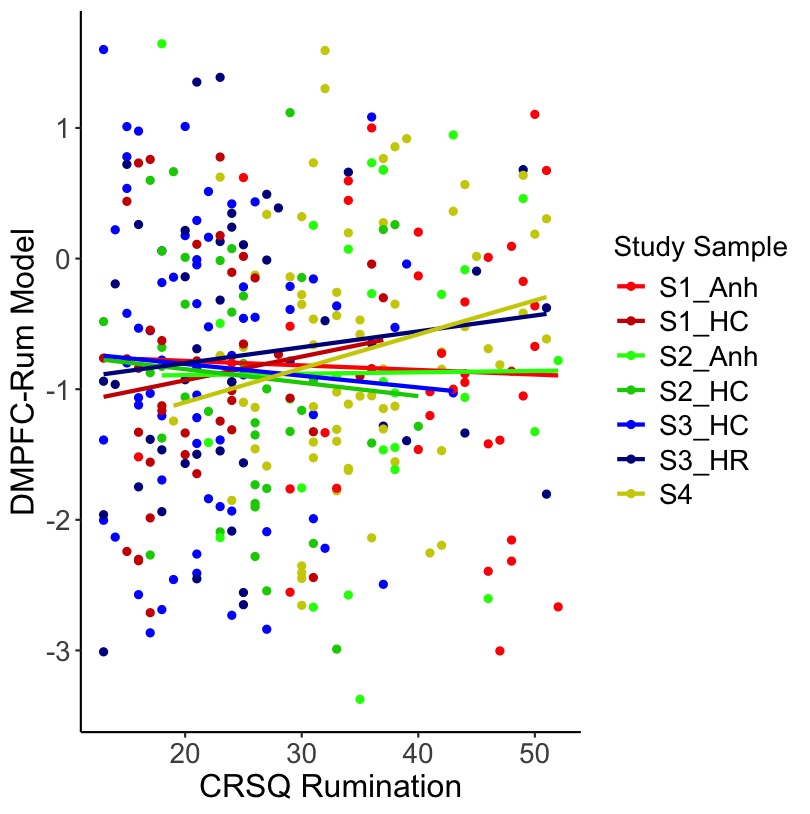

Figure 2: DMPFC-Rum model predictions of CRSQ rumination for internal data (Studies 1 to 4). Predictions based on DMPFC-Rum model from Kim et al. (y-axis), compared to actual CRSQ scores (x-axis). No consistent positive correlations are shown (see **Table S6** for stats). CRSQ: Child Response Style Questionnaire

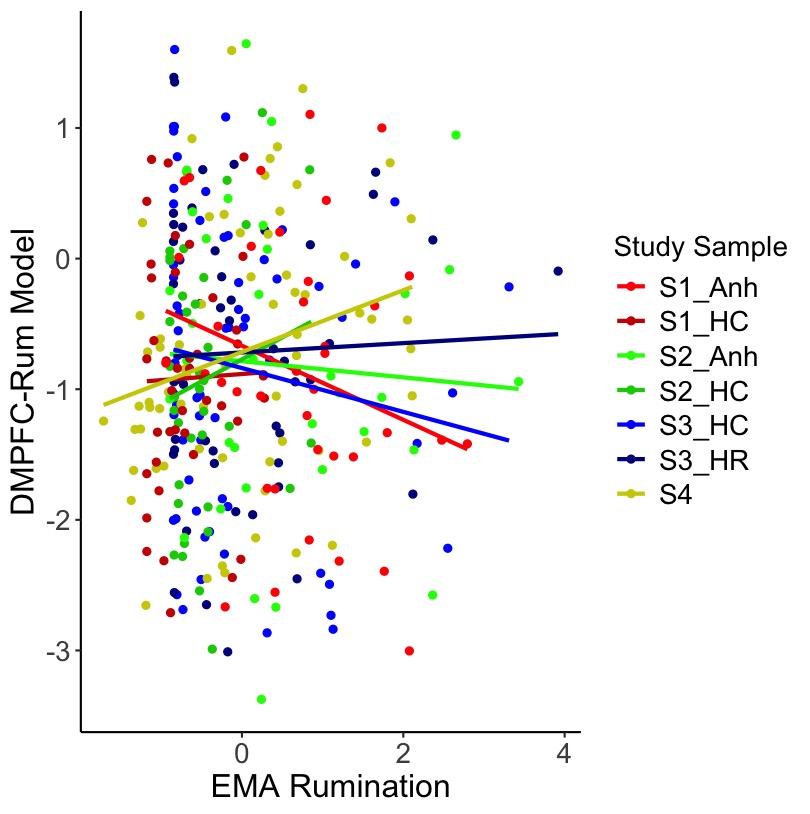

Figure 3: DMPFC-Rum model predictions of EMA rumination for internal data (Studies 1-4). Predictions based on DMPFC-Rum model from Kim et al. (y-axis), compared to actual EMA scores (x-axis). No consistent positive correlations are shown (see **Table S7** for stats). EMA: Ecological Momentary Assessment

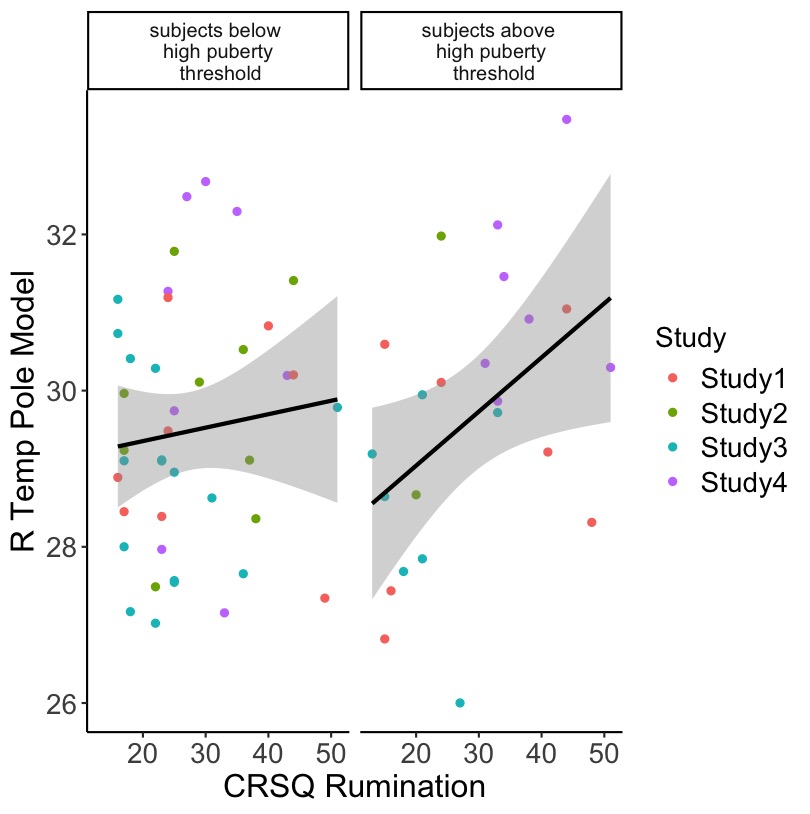

Figure 4: Right temporal pole model prediction interaction by puberty. On left, the relationship between right temporal pole predictions and actual CRSQ scores for low puberty. On right, the relationship between right temporal pole predictions and actual CRSQ scores for high puberty (0.5 * SD > mean).
